# Root responses to mechanical heterogeneity are shaped by pore geometry and root-soil biomechanics

**DOI:** 10.64898/2026.10.15.759134

**Authors:** Jiaojiao Yao, Gloria de las Heras, Daniel Patko, Jose Bico, Evelyne Kolb, Lionel Xavier Dupuy

**Affiliations:** PMMH, CNRS, ESPCI Paris, Université PSL, Sorbonne Université, Université Paris Cité, F-75005, Paris, France; Neiker, Department of Conservation of Natural Resources, Neiker, Derio 48160, Spain; Ikerbasque, Basque Foundation for Science, Bilbao 48009, Spain

## Abstract

Mechanical limitations to root growth occur when pressure exerted by turgor within root cells is counteracted by excessive resistance of the surrounding soil as well as by that of the cell walls. In a natural soil, however, heterogeneity allows roots to find paths of least resistance, though the governing principles behind this process have yet to be identified. Here, we combined artificial substrates, high-throughput live imaging, advanced image analysis, and mechanical modelling to uncover the principles governing the ability of a root to penetrate or deflect from zones of high mechanical impedance within a heterogeneous substrate. Our findings show that root responses to substrate heterogeneity can be accurately predicted from the pore-matrix geometry when both the mechanical properties of the substrate and those of the roots are known. Consequently, a crucial step toward understanding the physical limitations to root growth may lie in a detailed characterisation of root and soil micromechanical properties.

## Introduction

Soil structure, the way soil particles bind into macroscopic aggregates of various size and strength, is a key factor in the diagnosis of the quality of a soil and its ability to support a crop. It is often associated with organic matter content, water infiltration, nutrient availability and nutrient leaching (Bronick and Lal, 2005). However, soil structure is also a major factor in explaining the physical limitations to the growth of plant roots, a phenomenon exacerbated by increasingly heavy agricultural machinery (Keller and Or, 2022) and the rising frequency of droughts (Padrón et al., 2020).

Existing theories that link the mechanical properties of soils to root elongation represent the soil as a continuum, counteracting the growth forces generated by turgor pressure (Dexter, 1987; Tomobe *et al*., 2023; Koren *et al*., 2024). However, representing soil as a continuum is an oversimplification. Plant roots can exploit heterogeneities within the soil to find paths of least resistance (Popova *et al*., 2016; Martins *et al*., 2020; Wendel *et al*., 2022), which may explain why roots are observed to grow in soils where penetration resistance exceeds turgor pressure (Bengough and Mullins, 1990; Clark and Barraclough, 1999; Jin et al., 2013; Moraes et al., 2020). Numerous studies have described how roots interact with an obstacle through bending, buckling, thigmotropism and gravitropism (Whiteley *et al*., 1982). More recently, research has focused on understanding how roots perceive and respond to obstacles (Massa and Gilroy, 2003a,b; Silverberg et al., 2012; Jin et al., 2013; Bizet et al., 2016), and models are now able to explain root tropisms with a high level of detail (Porat *et al*., 2023). Nevertheless, applying these findings to identify the laws governing growth in a heterogeneous environment remains challenging.

To investigate root growth dynamics in heterogeneous substrates, we designed a simplified two-dimensional structured medium comprising agarose cylinders of different sizes and mechanical strengths, mimicking soil aggregates that roots may either penetrate or circumvent. The substrate also allows characterisation of the mechanical state of the system because properties of agarose can be obtained through mechanical tests, and the forces applied on the root through the contact geometry of adjacent cylinders. A high-throughput live imaging platform was employed to capture root trajectories with high accuracy, and a custom-made image analysis pipeline generated a database of root responses. Biomechanical models were then proposed to predict how factors such as pore geometry and the strength and stiffness of the gel influence root trajectories in heterogeneous media. Together, the results obtained demonstrate the potential to assess root growth based on the knowledge of root biomechanics and substrate heterogeneity.

## Material and Methods

### Fabrication of microcosm chambers

The microcosm chambers for plant growth were assembled from two parallel glass slides (76mm × 26mm × 1 mm, VWR, Spain) spaced with polydimethylsiloxane joins (PDMS SYLGARD^TM^ 184, Sigma-Aldrich, Spain). The PDMS U-shape part (Figure S1C) had a 3 mm by 3 mm cross section and allowed gas exchange and aeration of the sample (Liu *et al*., 2021). The two glass slides and PDMS were bonded by activating the PDMS surface via plasma treatment. Due to this process, the final microcosm chamber thickness was 2.7 mm, which is 0.3 mm smaller than the thickness of the initial PDMS U-shaped structure. Thus, the resulting volume of the microcosm chamber for root growth was 63.3 mm ×18 mm × 2.7 mm (respectively height, width, thickness).

### Preparation of the granular substrate

To facilitate the preparation of samples and analysis of root responses, we developed a two-dimensional granular substrate consisting of agarose cylinders. Agarose (BP160-500 lot 220906, Fisher bioreagents, Spain) powder was dissolved in deionized water at concentration of w/w, 1.5 %, 3.0 % and 6.0 % and subsequently heated at a temperature of 90°C until becoming transparent. The agarose solution was then poured into 3D-printed circular moulds (Figure S1A) with a thickness of 2.8 mm and a diameter ranging from 3.0 mm to 5.0 mm. The bottom of the mould was sealed with adhesive tape before pouring the agarose solution on the top of the mould. We allowed the excess of agarose solution to overflow the holes before sweeping the top of the mould. Once the gel had solidified, the cylinders were removed from the mould and collected in a Petri dish using a custom-made comb with cylindrical teeth matching the holes (Figure S1B). The initial thickness of the cylinders was 0.1 mm larger than the microcosm chamber thickness to ensure there would be no gap between the agarose cylinders and the microcosm chamber glass after drying and rehydration of the cylinders (see below).

To insert the agarose cylinders into the microcosm chambers, they were first oven-dried at 50 °C with air circulation until their thickness was reduced to 2.6 mm or less. Once outside the oven, deionized water was added on the surface of the cylinders to decrease friction with glass slides during introduction into the microcosm chambers. The cylinders were gently pushed and arranged in a triangular close packing within the microcosm chambers with a spatula. Finally, deionized water was introduced into the microcosm chamber, completely immersing the cylinders. The water allowed the cylinders to swell to a thickness that matches that of the microcosm chamber.

### Plant material

*Brassica oleracea* (cabbage) seeds (Corazon de buey pequno-balon, Fito, Spain) were soaked in deionized water at room temperature for 24 h. Seeds were then incubated in moist paper at 30°C during 24 hours and subsequently grown at 20 °C for another 24 hours. Radicles with length ranging from 4 to 6 mm were transferred to microcosm chambers for live imaging in high-throughput root phenotyping platform. The radicle was carefully inserted into the gap between two cylinders at the centre of the first layer. Then excess water surrounding the seed was removed using a dry tissue to ensure adequate aeration of the seed. Finally, parafilm tape was used to seal the microcosm chamber and to prevent the agarose gel from drying (Figure S1C).

### High-throughput root phenotyping platform and microcosm chamber fabrication

The high-throughput imaging platform (Figure 1A) consists of a large circular stage of radius 278 mm with slots for placing 25 microcosm chambers. The imaging system is fixed and consists of a Retiga R6 CCD sensor (Teledyn, UK) collecting 16-bit images formed by a large aperture imaging axis for large field of view (1X, 5.08 cm back aperture, 70 mm working distance, 20 mm field of view). Light was transmitted through the sample using a 3.2” TFT LCD panel, (1743 Adafruit, Mouser Electronics, Spain). The LCD panel was turned on only during image acquisition.

**Figure 1.**
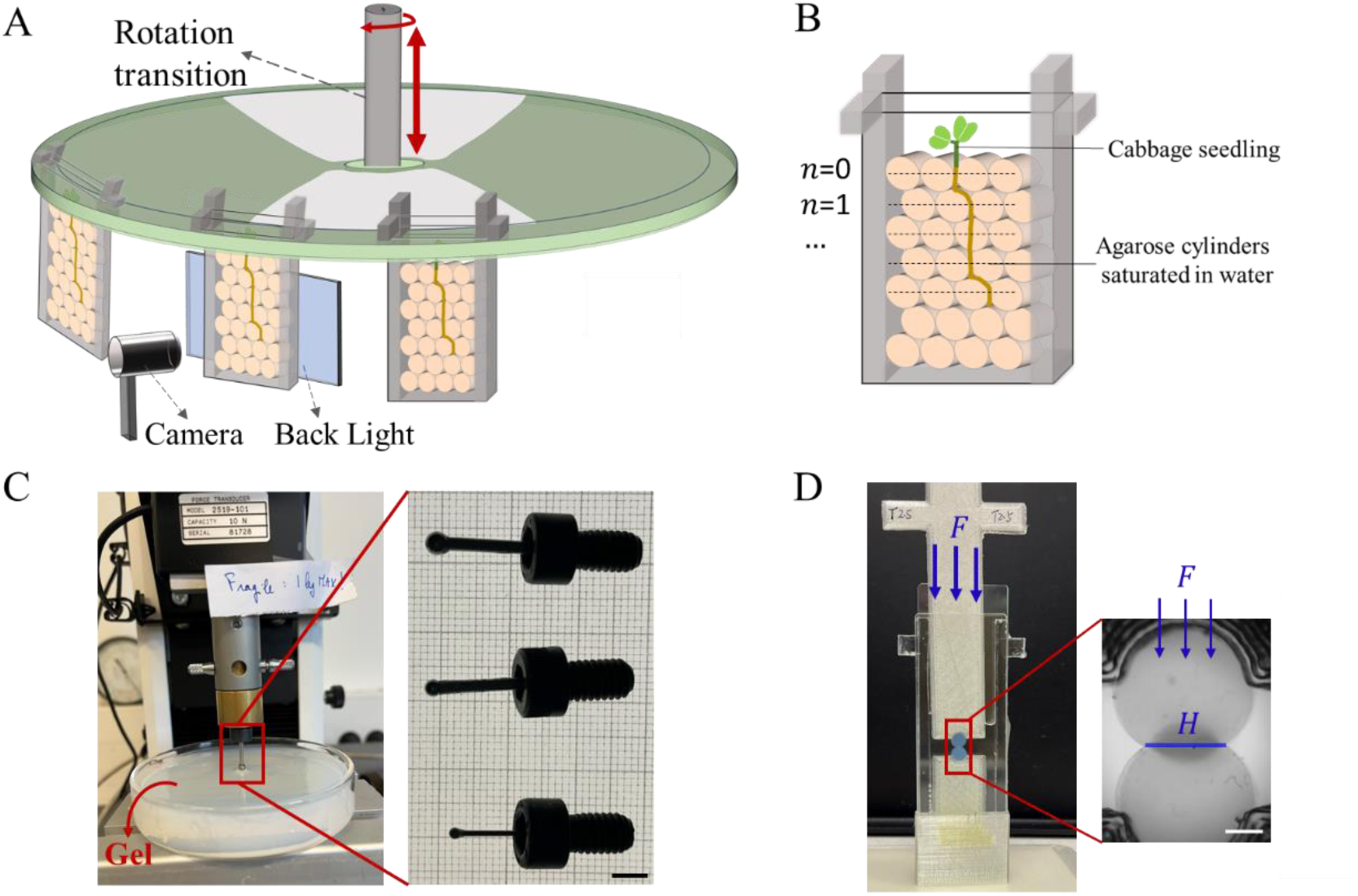
Experimental system made of gel cylinders of various size and strength. (A) A high throughput live imaging platform is used to capture time-lapse images of roots growing in heterogeneous substrates. Up to 25 microcosm chambers can be placed on a motorised stage in one experiment. A camera with a large field of view objective captures 6 images from each microcosm chamber. (B) The growth medium consists of gel cylinders of diameters 3.0, 4.0 or 5.0 mm with agarose gel concentration of 1.5%, 3% or 6%. The cylinders are arranged in a triangular close packing (Figure S1C). The internal thickness of the microcosm chamber is 2.7 mm and the initial thickness of the cylinder was adjusted to 2.8mm. (C) Indentation test used to determine the rigidity and strength of agarose gels. A universal testing machine (3340 Series universal testing systems, Instron, USA) with a force transducer of 10 N was used to establish the relationship between the loading force and indentation depth (left). Spherical indenters of diameters 1.6, 2.4 and 3.2 mm made of alloy steel were attached to a bolt for attachment on the load cell (right). (D) Compression test of agarose gel cylinders for the simultaneous acquisition of compression force *F_H_* and compression length *H*. Close up view of the system for positioning the agarose gel cylinders (left) and example image collected (right).

The sample stage has two degrees of freedom. Rotation is performed using a 2-inch aperture rotary stage (8MR190-2, Standa, Lithuania) and allows moving the microcosm chambers in front of the objective. Vertical movement of the microcosm chambers is performed using a motorized vertical translation stage (8MVT188-20, Standa, Lithuania) and allows acquisition of images from the same sample at different depths. The control of motion, illumination and image acquisition is performed using a custom-made program coded in Python (version 3.8). Six images corresponding to three vertical positions and two horizontal positions were acquired for each microcosm chamber at each timepoint.

### Experimental design

A fractional factorial design without interactions between factors was used to study the effects of agarose concentration and cylinder size. Microcosm chambers containing agarose cylinders of diameter 4.0 mm made from 3.0% agarose were considered as the reference condition and only one factor differed from the reference microcosm chamber in other microcosm chambers (Figure 1B). A total of 5 different conditions were studied, 3.0 mm diameter with 3% agarose, 4.0 mm diameter with 3% agarose, 5.0 mm diameter with 3% agarose, 4.0 mm diameter with 1.5% agarose and 4.0 mm diameter with 6% agarose.

Nine experimental runs with on average 15 microcosm chambers were performed with image data for a given microcosm chamber acquired every 30 minutes during 90 hours. For each run, two microcosm chambers were filled with plain agarose gels at each of the concentration found in other samples and used without seedlings as background sample for image processing. In the other microcosm chambers, only roots having grown continuously were kept for analysis, and about 30% of the samples were discarded because of slow or interrupted growth. Another 30% of the image data was not suitable for image analysis (presence of bubbles or roots out of focus). This resulted in a dataset comprising of 52 microcosm chambers with on average 10 replicates for each treatment but only four replicates when the diameter was 5.0 mm. Experiments were performed at room temperature monitored every 5 minutes throughout the duration of experiment (20 ± 3^°^C).

We also performed separate measurements of the root elongation rate of seedlings grown in microcosm chambers filled with plain agarose gels. The microcosm chambers were prepared as described above. 10 microcosm chambers were prepared with agarose gel of concentration 1.5%, 3%, 4.5% and 6%. After one day, we selected five microcosm chambers in which the roots had grown to roughly 10 mm in length. The microcosm chambers were subsequently transferred to the high-throughput imaging platform. Elongation rates were quantified by acquiring images immediately upon placement and one hour later.

### Image processing

Raw images were encoded in a 16-bit grayscale intensity values *I*_0_. To remove illumination artifacts, the background images *I_B_* (obtained from microcosm chambers filled only with agarose) were used to correct variations in intensities from the imaging system (flat field correction Figure S2 A&B),

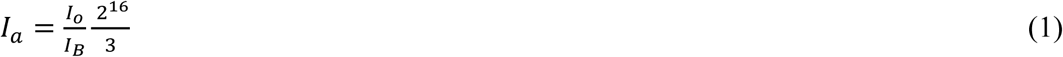

*I_a_* is the grayscale intensity image after flat field correction (Figure S2C). In this formula, the division is applied pixel-wise, and the factor 3 in the right-hand side is chosen arbitrarily to avoid spurious pixels exceeding the bit depth. The six images corresponding to a microcosm chamber were stitched into an image of approximately 5300 × 6100 pixels in size (Figure S2D) by copying them sequentially onto a regular array (left to right, top to bottom). No fusion step was required after flat-field correction, as each new tile simply overwrote the overlapping pixels.

A threshold and Euclidean distance transform were applied to the stitched images as a preparatory step for the determination of the centres of cylinders (Figure S3A). A watershed transform (Figure S3B) then segmented the cylinders from the images and centres were obtained as the centre of mass of each identified region (Figure S3C).

The stitched images were also processed using Kymolite, an application programmed in Matlab and adapted from Kymorod (Bastien *et al*., 2016). First a binary image of the root was obtained by thresholding. A contour detection algorithm was applied to extract the root centreline, which was subsequently used to calculate key variables such as root length, root tip angle, and incidence angles.

### Extraction of geometrical features from image data

The edges of the agarose gel cylinders were obtained by tracing rays from the centres of cylinders (Figure S4B) and identifying the maximum in the gradient of the pixel intensity along each ray (Figure S4C&D). This process was repeated for every one-degree rotation of the ray (360 rays traced). When two cylinders were adjacent, two pairs of nearest boundary points were identified (Figure S4E and inset) and the compression length (*H*) (Figure S4A) was then defined as the distance between the middle positions of these two pairs of points (Figure S4F). The compression length *H* was computed for each pair of adjacent cylinders located on the root path. The experimental determination of this length is a way to measure the compression force *F_H_* between cylinders.

The free length *L* is a proxy for the length of the segment of the root tip occupying the pore space (Figure S4 A). It was calculated as the distance between the end point of the compression length segment (blue asterisk in Figure S4 F) and the point where the vertical line passing through that location intersected the next cylinder (red dot in Figure S4 F). The incidence angle *θ_i_* is the angle between the root tip axis and the normal vector of the cylinder at the point of contact (Figure S5). The vector indicating the direction of the root tip was calculated by fitting a line to the last 0.25 mm of root centreline (identified as the length of rapid tapering of the root). The tangential direction was obtained by least-squares fitting a line to the boundary points corresponding to an arclength of 0.25 mm.

### Imaging of gel cylinders after penetration by plant roots

After completion of live imaging experiments, cylinders were removed from the microcosm chambers for further analysis. Cylinders that had not been in contact with or had been bypassed by the root were discarded. The cylinders which were penetrated by the root were kept for further analysis. A pseudo-Schiff staining protocol (Supplementary Method, section 1) was applied to bind propidium iodide to the free di-OH groups of agarose (Haseloff, 2003). In short, the agarose gel cylinders were first oxidized in a periodic acid solution for 30 mins. After rinsing, propidium iodide and Schiff reagent solutions were used to stain the agarose gel for 2 hours. The agarose gel cylinders were then rinsed and placed in a Light Sheet Fluorescence microscope (Liu *et al*., 2021). Images were acquired with an excitation wavelength of 514 nm and a 550 nm long pass emission filter (FGL550M, Thorlabs, UK).

### Characterisation of gel rigidity and strength

We used a universal testing machine (3340 Series, Instron) equipped with a 10 N load cell to characterise the Young’s modulus (*E_g_*) and the failure stress of the agarose gel (*σ_g_*). Spherical indenters of diameter of 1/16 inch, 3/32 inch and 1/8 inch (1.6 mm, 2.4 mm and 3.2 mm respectively, Figure 1C right) were adapted to the translation stage of the mechanical testing machine. Agarose gels were prepared at 1.5 %, 3.0 % and 6.0 % concentration and poured in a 90 mm × 20 mm (diameter × height) Petri dish in 18 mm-thick layers (Figure 1C left). A thin layer of deionized water was added on top of the gel to prevent it from drying and shrinking during the experiment. The test consisted in moving the indenter into the agarose gel layer and recording the relationship between the displacement of the indenter and the corresponding mechanical force resisted by the displacement of the indenter. The indenter was displaced at velocities of 0.01 mm s^−1^, 0.003 mm s^−1^, 0.001 mm s^−1^ or 0.0006 mm s^−1^, and the test was stopped shortly after the force ceased to increase due to the failure of the gel. The maximum force recorded was then used to determine the failure stress of the gel based on Hertz’ theory (Hu *et al*., 2010). Only a few trials could be achieved at a velocity 0.0006 mm s^−1^ due to software limitations and the excessive duration of such experiments.

### Characterization of the relationship between compression length and compression force

An additional mechanical test was performed to characterise the contact between adjacent cylinders. Two agarose cylinders were aligned vertically in a microcosm chamber using 3D printed holders (Figure 1D). The top cylinder was moved at a speed of 0.017 mm/s using a lifting stage. A precision scale (ME 4002, Metler Toledo) was used to record the increase in force resulting from the movement of the top cylinder against the bottom cylinder (Figure S6A). Images were acquired at one frame per second, using the light transmitted from the LCD panel. The acquisition of data from the precision scale was computer controlled to synchronise with image acquisition. The compression length between two agarose gel cylinders was measured from image data (Figure S7). Data was recorded only when the compression length began increasing steadily. The initial stage of the test was discarded because factors such as movements of the cylinders and sensor drift produced spurious data. We performed tests for cylinders of diameters 3.0 mm and 5.0 mm at 3% agarose gel concentration, and with diameter of 4.0 mm at 1.5%, 3% and 6.0% agarose gel concentration.

### Model Parameterisation

The compression length *H*, free length *L* and incidence angle *θ_i_* were extracted using the image analysis pipeline presented above. The Young’s modulus *E_g_* and failure stress *σ_g_* of the gel were determined from mechanical testing experiments. The root diameter was fixed to *D_r_* =0.3 mm based on measurements from image data. The Young’s modulus of the root was estimated to *E_r_* = 20 MPa based on values reported for the growth zone of Maize (Quiros, 2023).

### Software for data processing

All algorithms for image processing, image analysis, feature extraction or for computing model outputs were developed using Matlab 2022b (MathWorks). Kymolite, the modified version of Kymorod was also written in Matlab 2022b.

## Results

### Mechanical properties of agarose gel

Results of indentation of bulk agarose gel were first performed using different indenter sizes and displacement rates, and then analysed using formulas from Hertz contact to estimate the Young’s modulus and failure stress of the gel, *E_g_* and *σ_g_* (Supplementary Method, section 3, equations 7 and 10). Estimations were independent of the size of the indenter which confirmed the validity of the test. Given that root elongation occurs at approximately 10^−4^ mm s⁻¹, we performed the indentation tests on bulk agarose gel using the lowest displacement rates achievable.

Young’s modulus and failure stress obtained at displacement rates in the range 6 × 10^−4^ to 3 × 10^−3^ mm s^−1^ (Figures 2A&C) were comparable, and we chose to characterise the different agarose gel concentrations using displacement rates of 3 × 10^−3^ mm s^−1^. Although this shortened the overall duration of experiments, each test still required more than one hour to be completed. Experimental data showed the failure stress *σ_g_* and the Young’s modulus *E_g_* of the gel had similar values for all agarose gel concentrations (Figure 2C). Using these settings, the failure stress and Young’s modulus of the agarose gel varied linearly with agarose gel concentration *C_g_* in the range [1.5%, 6%], following the fitted equation *σ_g_* = 81.4 *C_g_* − 33.6, *R*^2^ = 0.99 (Figure 2B, red curve) and *E_g_* = 90.5*C_g_* − 67.7, *R*^2^ = 0.99 (Supplementary Method, Figure 1), with *σ_g_* and *E_g_* in kPa and the concentration of the agarose gel *C_g_* given in mass percentage.

**Figure 2.**
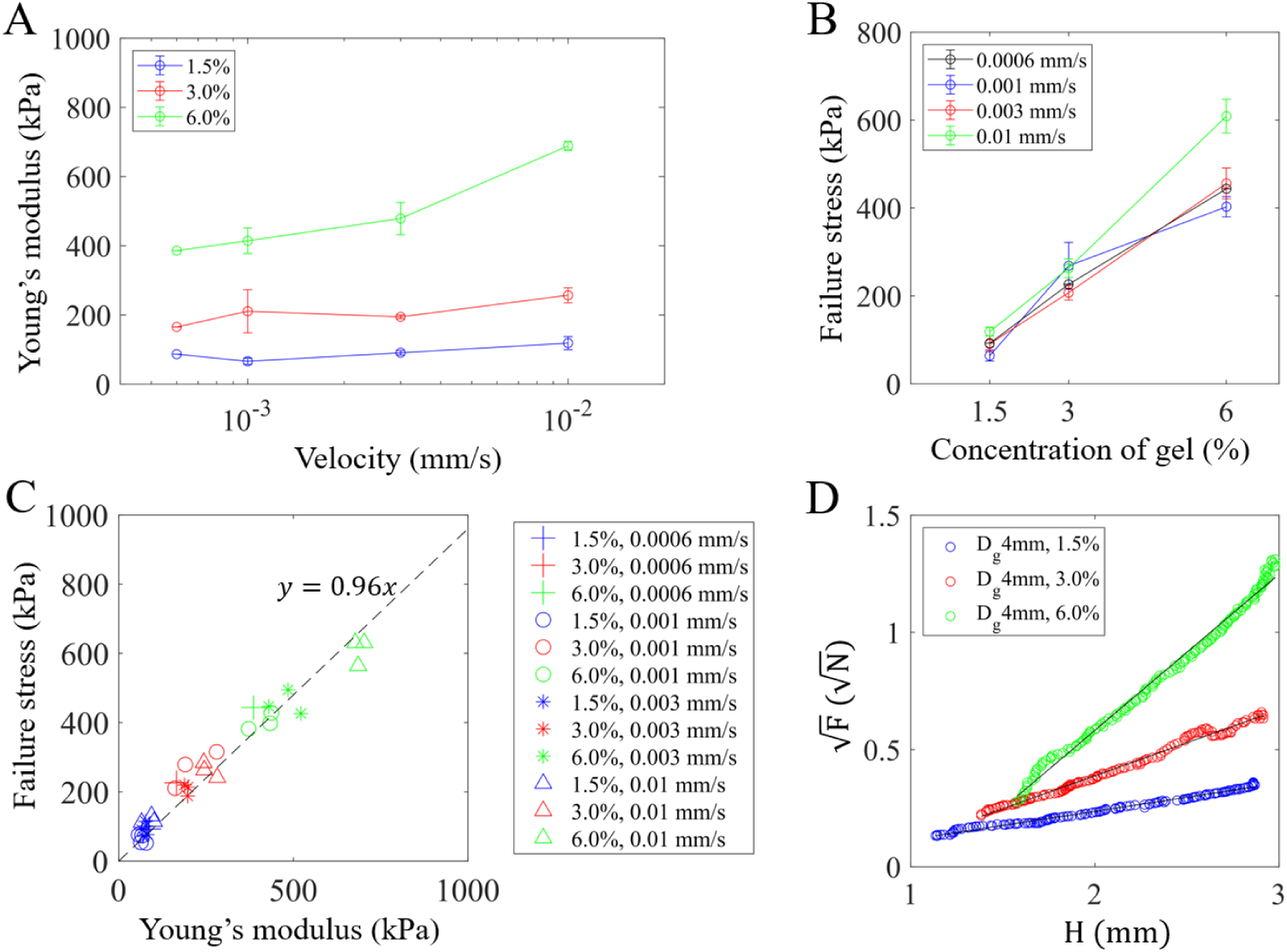
Mechanical properties of agarose gel and agarose gel cylinders. (A) Young’s modulus of agarose at different agarose gel concentration as a function of the velocity of indentation (here data shown for tests with 1.6 mm diameter indenter). (B) The failure stress varied linearly with the agarose gel concentration (here data shown for tests with 1.6 mm diameter indenter, colours indicate different indenter displacement velocities). (C) Linear relationship between the failure stress and the Young’s modulus of the agarose gels (the black dashed line represents the linear fit for the entire dataset). (D) Compression tests between cylinders of diameter of 4.0 mm showed that the relationship between the compression length and the square root of the force can be approximated with a linear relationship, which confirms compression of gel cylinders follows the Hertz contact formula (Supplementary Method, section 2).

The compression tests performed directly on agarose gel cylinders showed the compression length *H* is related to the square root of the measured force *F_H_* holding two cylinders in contact, which is in good agreement with Hertz contact theory (Figure 2D). Collectively, results in this section confirm that the model system can be characterised using classical solid mechanics and provides data valuable for the development of a biomechanical model (Supplementary Method, section 4).

### Root response is affected by the concentration and diameter of the agarose gel cylinders

Root growth in the absence of heterogeneity, that is in the bulk agarose, gel was influenced by the agarose gel concentration. The elongation rate observed was 0.43 ± 0.07 mm h^−1^ (n=5), 0.34 ± 0.11 mm h^−1^ (n=5), 0.19 ± 0.17 mm h^−1^ (n=4) for seedlings grown in agarose gel concentration of 1.5%, 3% and 4.5% respectively. Because roots did not grow in 6% plain agarose gel, we do not report elongation rates for this treatment. Root growth in the triangular close packing of agarose gel cylinders was investigated by decomposing the root’s trajectory into successive elementary events corresponding to the passage of the root tip from row (*n* − 1) to the next horizontal row (*n*) of cylinders. We studied only the cases where the basal part of the root was held between the two agarose gel cylinders of the row *n* − 1. This ensured that the forces confining the base of the root were well-defined (Figure 3A). Under these conditions, the root either penetrated the underlying agarose gel cylinder at row *n* (hereafter referred to as a ‘penetration’ event, Figure 3B middle) or deflected from it (hereafter referred to as a ‘deflection’ event) (Figure 3B left). In a few cases, the root tip remained pinned for extended periods of time at the top surface of the gel cylinder before generally deflecting. (Only in one case this behaviour led to a penetration event.) In these later cases, roots exhibited behaviour distinct from other events and were classified as ‘delay’ events (Figure 3B right). To confirm root growth did not disturb the packing or modified the forces holding the cylinders together away from the root path, we measured the mean compression length *H* in 5 microcosm chambers at the beginning and end of the experiments, excluding those compression zones along the path of the root. The difference in compression length was 0.12 ± 0.12 mm, and a two-sample t-test showed it was not significantly different from 0 (p=0.38, n=25). Consequently, the passage of the root between two cylinders does not alter significantly the compression forces existing between other couple of agarose cylinders. Therefore, the compression length (hence the compression force) was extracted from the first image acquired on a sample before the root grew in the microcosm chamber (Figure 3A).

**Figure 3:**
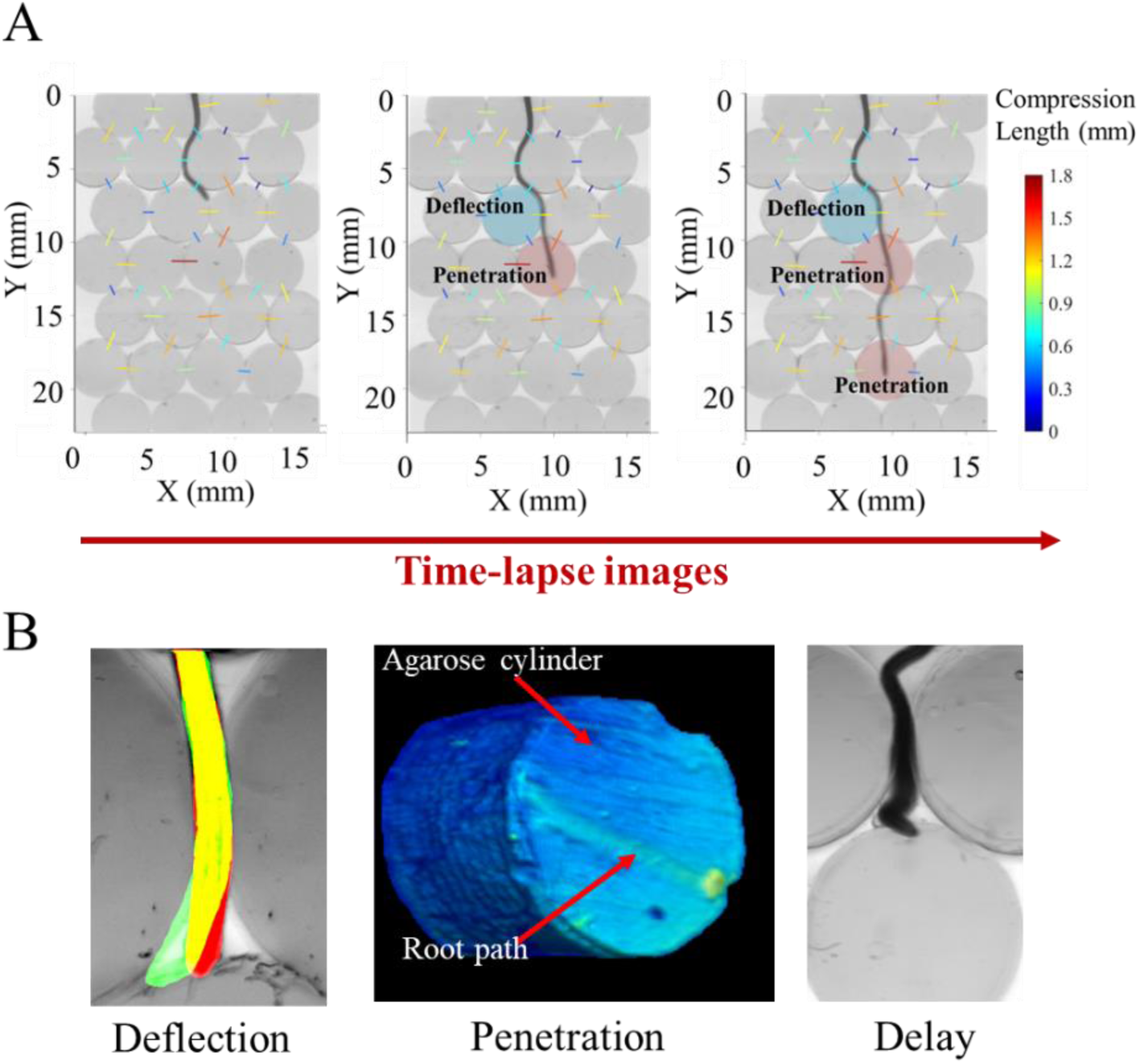
Root growth in a triangular close packing of agarose gel cylinders. (A) The trajectory of the root results from a series of penetration and deflection events depending on the pore geometry, the mechanical properties of the gel and the distribution of contact forces between agarose gel cylinders (presented here as compression length using a colour map, with the cylinder diameters of 4 mm). (B) The study focuses on the response of roots that are held at their base between the two lateral cylinders while growing toward a third cylinder positioned below, in front of the root tips. In these conditions, a root may either deflect (left, red indicates the position of the root at the moment of contact and the green indicates the position of the root following the deflection). The root may also penetrate the agarose cylinder (middle). In this case, light sheet microscopy data shows the root is able to damage the gel. In a few cases, the root tip became pinned in the agarose gel, and only after a long delay was the root eventually able to bypass the agarose cylinder. These cases were termed a delay event (right).

3D light sheet fluorescence microscopy data of the agarose gel cylinders used in experiments confirmed that root penetration events caused permanent damage to the gel, even when growing between the gel and the glass surface (Figure 3B, middle). This damage confirmed that the gel’s failure stress is a relevant mechanical property for characterising root penetration, directly affecting the ability of the root to penetrate the cylinders. Roots growing in substrates made of agarose gel cylinders of diameter 4.0 mm and concentration of 1.5%, 3.0% and 6.0% were able to penetrate the gel with a probability of 50.0%, 38.2% and 3.1% respectively (Figure 4A&B). Less intuitively, visual observation also indicated that the size of the agarose gel cylinders affected the ability of the root to penetrate the gel (Figure 4C). For example, roots growing in substrates which cylinders had diameter of 3.0 mm, 4.0 mm and 5.0 mm were able to penetrate the 3% agarose gel cylinders with probabilities of 70.6%, 30.2% and 25.0% respectively (Figure 4D).

**Figure 4.**
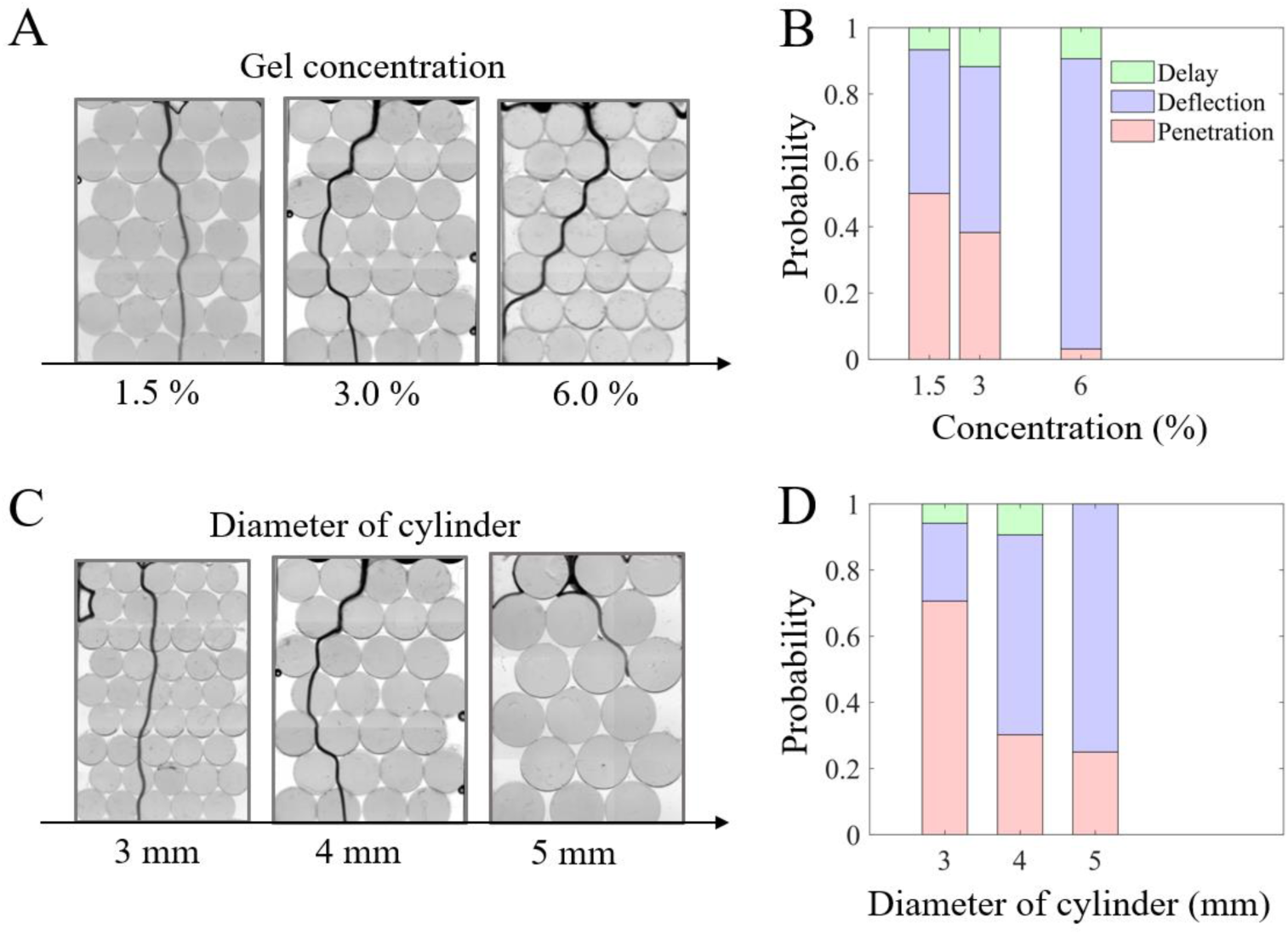
Root trajectories vary as a function of agarose gel concentration and cylinder size. (A) At constant diameter (4.0 mm shown here), the increase in agarose gel concentration leads to more root deflections and increased root tortuosity. (B) Probability of an event of penetration, deflection or delay to occur at each contact with an agarose gel cylinder of 4.0 mm diameter as a function of the agarose gel concentration. (C) The increase in diameter of the agarose gel cylinder at a given agarose gel concentration (3.0% shown here) also results in more root deflections and morphologies that appear more tortuous. (D) Probability of an event of penetration, deflection or delay to occur at each contact with a cylinder with an agarose gel concentration of 3.0% as a function of the diameter of the cylinder. Probabilities of a given event are calculated as number of events divided by total number of interactions between a root and a cylinder.

### Geometrical properties of the pore space

To explain these results, we then looked into the geometrical properties of the pore space resulting from the triangular close packing of the cylinders (triangular arrangement of the cylinders). Theoretically, the free length (*L*, also a proxy for the pore size) and therefore the ability of the root to deflect is directly related to the diameter *D_g_* of the agarose gel cylinders and the compression length *H*. A first order approximation in 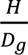 of the calculated free length is 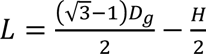, where 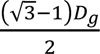 is the pore size when *H* = 0 in a triangular close packing of perfectly arranged and monodisperse cylinders. Experimental characterisation of the arrangement of the agarose gel cylinders confirmed the triangular close packing approximation, although variations around the expected values were observed (Figure 5A). Due to the free space in the pore size, the root had the possibility to deviate from a straight vertical growth. This induced variations in the angle of incidence at the time of contact with the next cylinder. Measured values of *θ_i_* reached up to 70° (Figure 5B) but the probability of finding a non-zero angle of incidence declined rapidly with the median angle characteristic of the distribution of 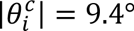.

**Figure 5:**
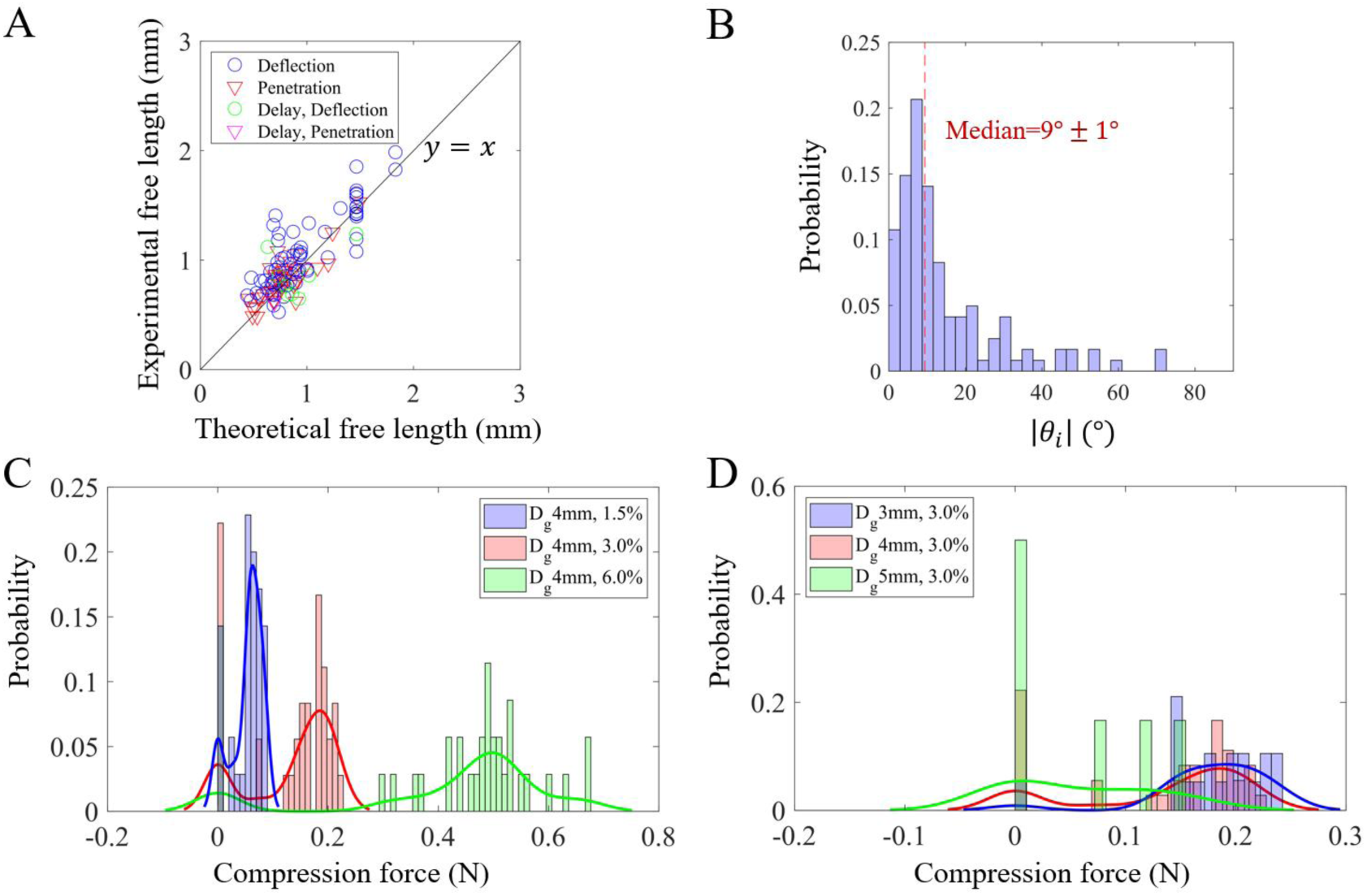
Characterisation of the model system. (A) Experimental free length plotted as a function of the calculated (theoretical) free length 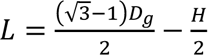. Colours indicate the root behaviours (red for penetration, blue for deflection, green for delay event before deflection, magenta for delay event before penetration). (B) Histogram of the frequency of the absolute value of incidence angle *θ_i_*. (C-D) Histogram showing the distribution of compression forces resulting from the contact between adjacent cylinders. Compression forces *F_H_* are calculated from the experimental determination of the compression length *H* through the formula 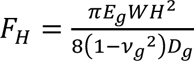 with *ν_g_* = 0.5 (C) for various agarose gel concentrations at constant diameter of cylinders (*D_g_* = 4.0 mm) (D) for various diameters *D_g_* of the cylinders at constant agarose gel concentration (3%). Legend captions for (C) and (D) indicate diameter and agarose gel concentration respectively.

The arrangement of the cylinders within the microcosm chambers was influenced by several factors. These included the swelling of the cylinders following water addition and the constraints imposed by the front and side walls, which limited expansion and generated shear along the walls and compression forces that held the cylinders in place. To characterise these constraints, we measured the compression lengths *H* and infer the mechanical forces holding the cylinders in place within the microcosm chambers. We plotted the distribution of compression forces holding cylinders along the path of the roots. We observed a bimodal distribution of the contact forces between the agarose gel cylinders. The first mode corresponded to cases where a cylinder was not in contact with its neighbour (*H* = 0), while the second mode was associated to cases where a finite pressure held two adjacent cylinders. There was an effect of the agarose gel concentration on the levels of compression forces (Figure 5C) but there was no effect of the size of the cylinder (Figure 5D). The swelling of agarose that occurs during substrate preparation may play an important role in the development of compressive forces between adjacent agarose gel cylinders. Because higher agarose concentrations produce stiffer gels, the confinement imposed by the microcosm chamber may increasingly restrict gel expansion during the swelling of the gel, and the constrained conditions may generate greater compressive forces between cylinders at higher gel concentrations.

### Simplified model for the penetration of a root

Different types of mechanical interactions occurred between the root and the surrounding agarose gel cylinders (Figure 6A, superimposed images obtained before and after the root overcame the resistance from the gel). We observed that (i) the roots could pass through without bending by penetrating the agarose gel cylinder; (ii) the root bending allowed the tip to slide past the agarose gel cylinder; (iii) the support provided by the gel cylinders surrounding the base of the root deformed during the bending of the root apex, which permitted further lateral displacement of the root tip; (iv) the support of the base of the root was kept in place and the pinning of the root tip in the agarose cylinder below, evidenced by visible deformation of the gel, constrained the lateral displacement of the tip. These boundary conditions resulted in a complex deformed root shapes above and below the compression zone. In some rare cases the fourth case led to a delay event.

**Figure 6:**
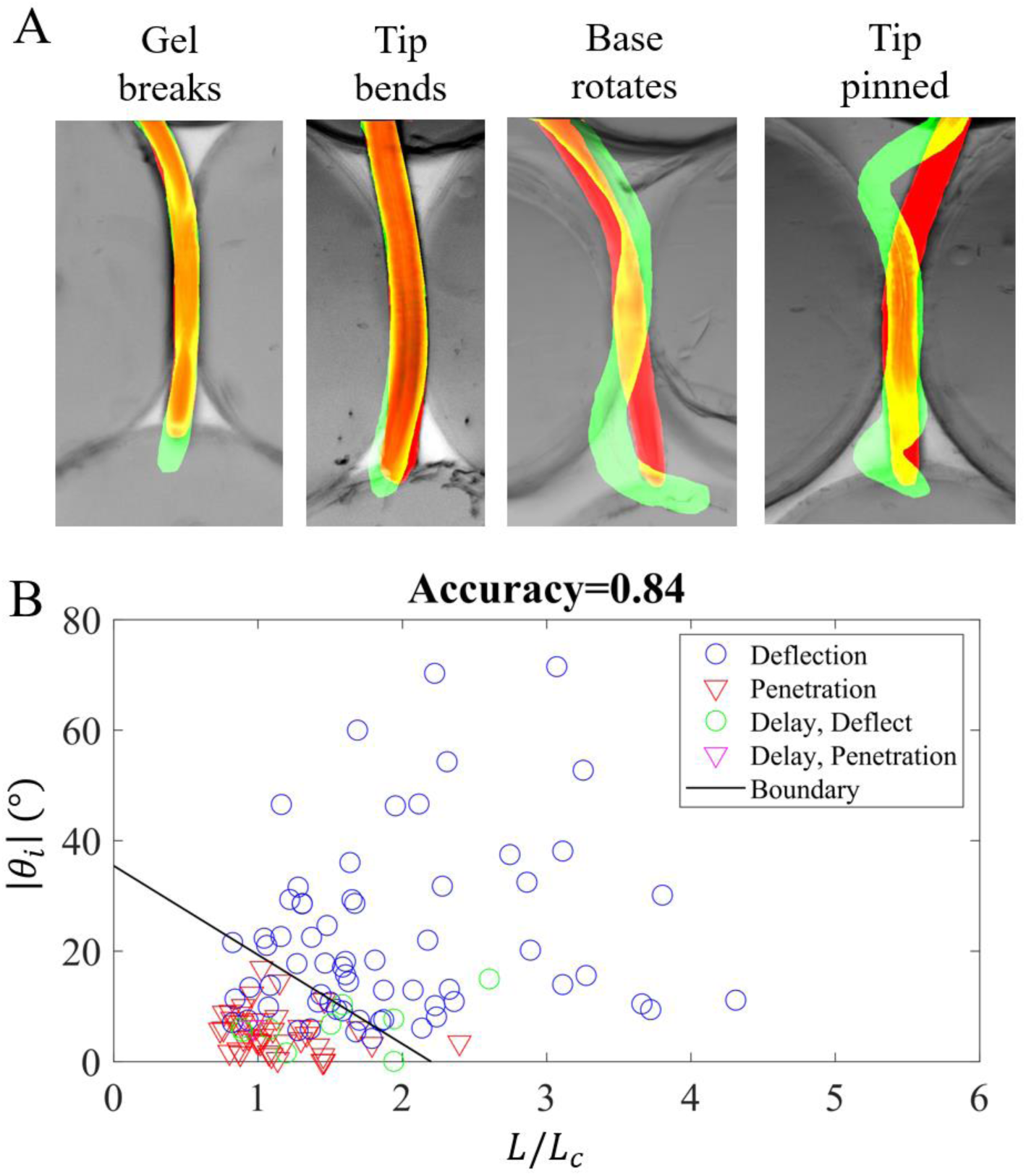
Possible interactions between a growing root tip and agarose gel cylinders. (A) Potential mechanisms leading to the failure of the root-gel system. Red and green colours indicate the position of the root before and after contact with the cylinder, respectively. The base is the part of the root squeezed between the two lateral cylinders. (B) Absolute value of the incidence angle versus the dimensionless length ratio 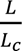, where *L* is the experimental free length and *L_c_* is the critical length, with 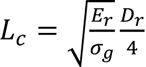. The straight line shows the boundary between deflection and penetration cases obtained from the linear discriminant analysis.

These observations led us to interpret root behaviour within a biomechanical framework. Following contact with the lower agarose cylinder, root elongation becomes axially constrained, causing the axial compressive force *F* experienced by the root to increase progressively. Due to Newton’s third law, the magnitude of this force equals the force exerted by the root on the lower cylinder. For the sake of simplicity, we assumed that the root axis in between the base and the tip has a constant diameter *D_r_* and is mechanically homogeneous with a uniform Young’s modulus *E_r_*. Under these assumptions, the root is expected to buckle if the axial compressive force reaches the classical buckling force 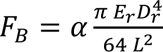 (Supplementary Method, section 4 equation 11) where *α* is a dimensionless numerical prefactor of order 1 that depends on clamping conditions at both ends of the root (base and tip). Alternatively, the root might penetrate the gel if the failure stress *σ_g_* of the gel is reached. We assumed that this happens when the axial force exerted by the root reaches 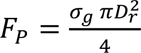 (Supplementary Method, section 4, equation 12). If the axial force *F* first reaches the penetration force of the gel *F_P_*, the root is able to penetrate the agarose gel cylinder. Conversely, if the critical buckling force *F_B_* is reached first, the root buckles and bends. Therefore, in the case where *F_P_* < *F_B_* the root is predicted to penetrate the agarose gel, while in the opposite case (*F_B_* < *F_P_*), the root is predicted to buckle. The criterion *F_B_* = *F_P_* marks the transition boundary between the two behaviours. Since *F_B_* depends on the free length (*L*), this criterion can be written in terms of a characteristic length 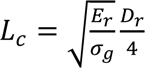. The length scale *L*_c_ only depends on the mechanical (*E_r_*) and geometric (*D_r_*) properties of the root and on the strength (*σ_g_*) of the surrounding gel. The criterion setting the transition from penetration to buckling can therefore be related to the size of the pores, for which *L* here serves as a proxi. We thus expect the root to penetrate for 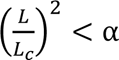 and conversely to buckle for 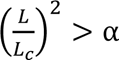.

However, the comparison of *L* to the characteristic length *L_c_* is not the sole criterion dictating the ability of the root to penetrate the gel. Next, we examined how the incidence angle *θ_i_* affected the ability of the root to penetrate agarose gel cylinders. The relationship between |*θi*|, the ratio 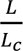 and the ability of the root to penetrate was studied using a linear discriminant analysis (Figure 6B). The analysis showed that, in the 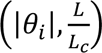 two-dimensional space, cases of deflection are clearly separated from cases of penetration. When the angle of incidence was zero, no penetration occurred for 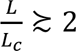, resulting in *α* ≈ 4. The order of magnitude of the pre-factor was in reasonable agreement with a mechanical buckling mechanism. Indeed, we would expect a value of 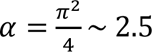 for the buckling of a rod fixed at one end (root base being blocked by the two lateral cylinders) and free to rotate and move laterally without any appreciable friction at the other end (root tip). Alternate scenarios involving pinned-pinned or fixed-pinned configurations would lead to higher values of *α*, of *π*^2^ and 2*π*^2^ respectively, or even 4*π*^2^ in the case of fixed-fixed configurations. For a given agarose gel concentration and diameter of cylinder, we could expect the value of *α* to increase with the lateral compression force maintaining the root base. However, drawing a definitive conclusion for the local nature of the clamping conditions would require precise experiments with a model system, which is beyond the scope of the present work.

As |*θ_i_*| increased (Figure 6B), the 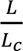 threshold at which roots could no longer penetrate decreases, indicating that the mechanical constraints preventing deflection weakened. This behaviour can be explained by the fact that the forces acting on the root are no longer purely axial. Almost no penetration was observed for incidence angles larger than 15°. This specific value may be interpreted in terms of Coulomb friction framework, where the tip of the root is expected to slide for an incidence angle larger than 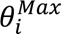(this would lead to a friction coefficient 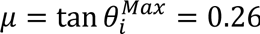). Incidentally, value of 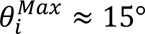 is of the same order of magnitude as the characteristic angle of the distribution of incidence angle of the data, (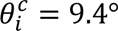, Figure 5B). However, the friction of a root on a soft hydrogel may not follow classical Coulomb friction. The detailed nature of the interaction remains an open question.

### The ability to penetrate soft obstacles affect the efficiency of root system

The capacity of roots to deflect can represent both a benefit and a limitation for the plant. Deflection enables roots to grow along the zones of least resistance in the soil. However, it also makes their trajectories more tortuous, resulting in the root system spreading horizontally rather than penetrating deeply. To analyse this process, we grouped image data according to agarose gel concentrations (Figure 7A) and diameters of the cylinder (Figure 7B). The root skeletons of the seedlings from different replicates were laterally translated to align their planting positions before superimposition. Observations confirmed a relationship between the lateral spread of the root skeletons and the probability of deflection. At high agarose gel concentration (6.0%), the penetration of agarose gel cylinders was rare (about 3.1% probability) resulting in tortuous root paths. As agarose gel concentration decreased, roots were more vertical and straighter, due to the increased probability of penetration. The effect of the diameter of the cylinder was less clear due to a lack of data for *D_g_* = 5 mm (less root cylinder interactions due to a greater number of roots reaching the side wall).

**Figure 7.**
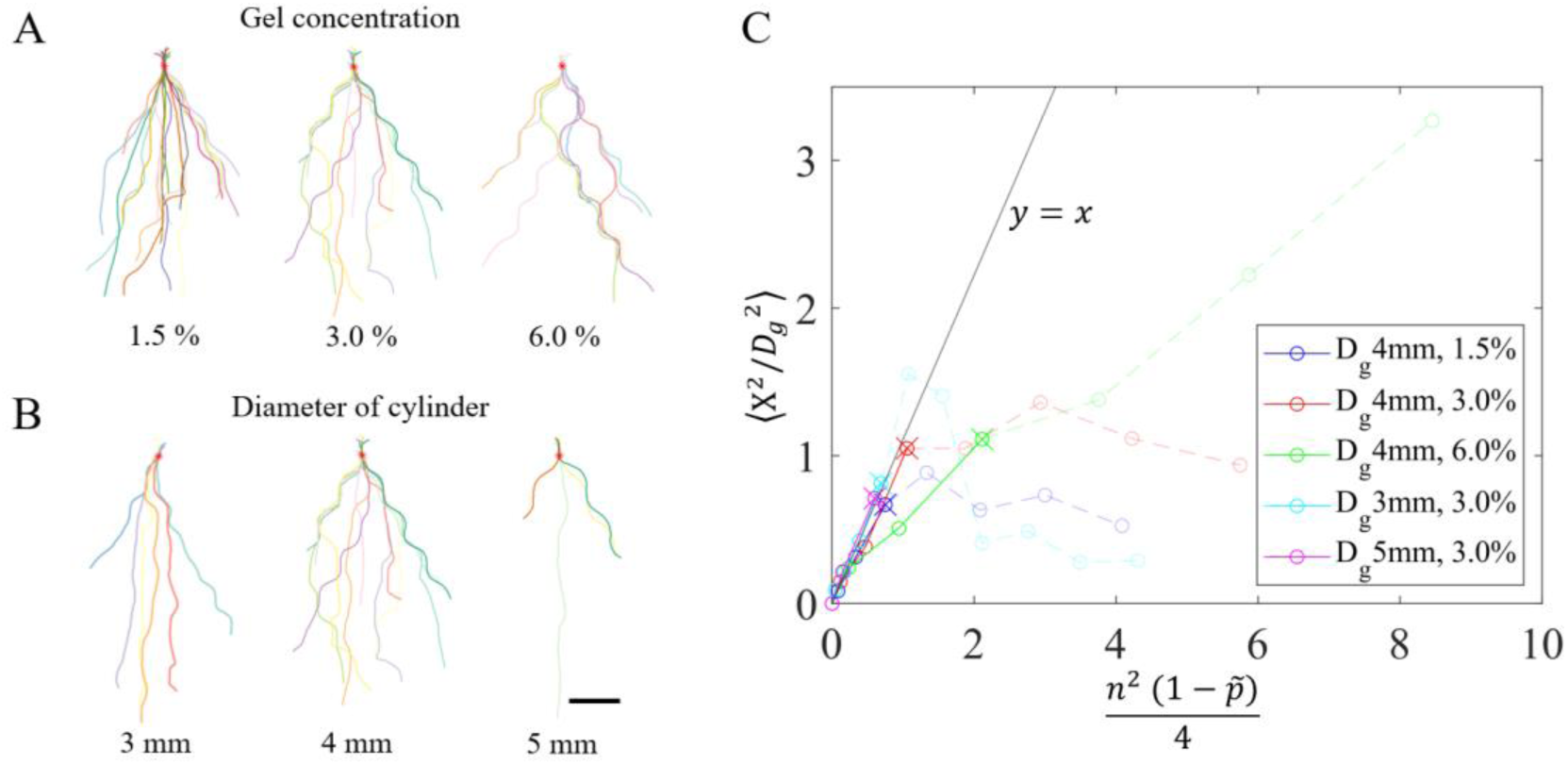
Effect of substrate composition on rooting’s lateral spread. (A) Root skeletons in substrates of increasing agarose gel concentration (4 mm cylinders’ diameter at respectively 1.5%, 3.0% and 6.0% agarose gel concentration). (B) Root skeletons in substrates of increasing cylinders’ diameter (diameters of respectively 3.0 mm, 4.0 mm, and 5.0 mm at 3.0% agarose gel concentration). The red asterisks mark the origin, which is defined as the point where the planting position of the seedlings intersects with the horizontal centerline of the first row. Scale bar represent a length of 5 mm. (C) Evolution of the dimensionless variance of the lateral spreading (root tip’s horizontal distance from the origin) with a normalised variable depending on the row number *n*. Details of the derivation of these variables are provided in equation 20 of the Supplementary Methods. The straight line *y* = *x* represents the relationship predicted by the biased random walk model.

To analyse the root trajectory data within a theoretical framework, we first constructed a simple random walk model. The model relates the horizontal position of the root tip *x* to the row number *n* and to growth and morphological variables such as the horizontal spread of the root tips, total length of the root or the time needed to reach the row *n*. The model assumes that the root elongation rate in the gel *v_g_* is constant and slower than the elongation rate in the pore *v*_0_ which is also assumed to be constant. We defined the global probability 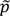 that a root is growing straight from row *n* to *n* + 1(Supplementary Method, section 5). If the root deflects to the right and to the left with the same probability 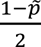, the expected variance of the horizontal position *x* of the root tip increases linearly with *n* after going through *n* rows of cylinders and is expressed as 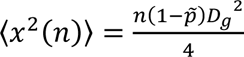 (Supplementary Method, equation 18). Using the probabilities of penetration obtained experimentally (Figure 4), the model consistently underestimated the lateral spreading of the roots ⟨*x*^2^(*n*)⟩ observed experimentally (Supplementary Method, section 5, figure 4).

Next, we hypothesised that the underestimation of the random walk model was due to the directional bias in root deflections. Indeed, most roots exhibited oblique trajectories (Figure 7A&B) whereas a random walk with symmetric probabilities would be expected to generate numerous path crossings. The model was therefore modified to represent the following limit case: the root either penetrates or deflects toward a given direction, that is the direction of deflection is conserved for the oblique parts of the trajectory. In this case, the expected horizontal spread of the root is 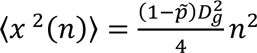 (Supplementary Method, section 5, equation 20). Using this model, the data from the various agarose gel concentrations collapsed into a single relationship matching well the model predictions when growing through the first few rows of agarose gel cylinders (Figure 7C). Beyond a certain depth, roots started reaching the side of the microcosm chambers (indicated as a cross in Figure 7C) and were subsequently discarded from the analysis. The experimental variance beyond that depth is that of a probability distribution truncated by the boundaries of the microcosm chamber and therefore the variance was logically overestimated by the model. Also the possibility switching between right and left deflections was not taken into account, which would also lead to model overestimation, see for example root growth in 4 mm gel cylinders at 6% agarose gel concentration (Figure 7C).

Importantly, the interest of the random walk model was to highlight the key physical parameters for describing the growth of root systems in heterogeneous media. First, in the simple random walk model, as expected, the increased global probability of growing through gel (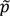) reduces the length of roots required to reach a given depth, 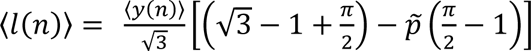. However, this feature comes with a potentially increased time needed to reach this depth at row *n*. The corresponding time 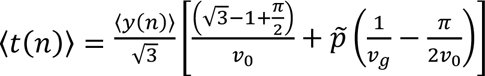 in this case is dependent on the root elongation rate in the gel *v_g_* and on the unconstrained root elongation rate *v*_0_ inside the porous network. This time increases with 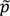, unless the elongation rate *v_g_* in the agarose gel is large enough such that 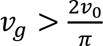. Based on the root elongation rates measured in 1.5%, 3%, and 4.5% agarose gels (see earlier section “Root response is affected by the concentration and diameter of the agarose gel cylinders”), higher probability of deflection (smaller 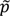) would decrease the duration to reach a given depth, hence decrease the velocity of the rooting front at gel concentrations above 3%. Finally, if we define the length to depth ratio 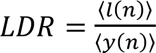 where *l*(*n*) is the length of the root at row *n* and *y*(*n*) is the corresponding depth at row *n*, then the inverse of *LDR* is an indication of how efficiently the root front progresses vertically and the random walk models predict that the *LDR* is only affected by the structure of the substrate through the probability of penetration 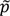.

## Discussion

Root system architecture is a key determinant of crop productivity (Rogers and Benfey, 2015), yet predicting its development from knowledge of soil conditions remains a major challenge. The lack of fundamental laws describing how physical constraints on growth, especially those arising from soil heterogeneity, limits our ability to manage crops efficiently for reduced input or enhanced abiotic stress tolerance. Results in this study support the idea that reductionist approaches can be employed to predict the influence of substrate structure on the spread of the root system. Several properties likely govern the ability of the root to maintain stable growth in structured soil. The strength of the matrix material (here *σ_g_*) is critical factor affecting root growth. An increased ratio between the mechanical properties of the matrix *σ_g_* and those of the root *E_r_* increased the likelihood of deflections and therefore the tortuosity of the root. Growth mechanisms in structured soil are therefore not uniquely associated with the resistance from the medium to elongation and therefore differ from the classical frameworks for predicting root elongation as a function of external pressure (Bengough and Mullins, 1991; Cosgrove, 2016; Trinh *et al*., 2021).

The size and shapes of pores can vary very significantly between soils, as a function of soil texture (Rabot *et al*., 2018), aggregation (Pagliai and Vignozzi, 2002), organic matter content (Tisdall and Oades, 1982; Pagliai *et al*., 2004), or soil management (Pagliai *et al*., 2004). In our experiments, the pore size was controlled through the diameter of agarose gel cylinders (*D_g_*), but it also varied stochastically within a sample due to differences in the compression forces holding the cylinders within the microcosms, with broad distributions reminiscent of those observed in a granular medium (Fakih *et al*., 2019). The dominant effect of increased pore size is on the free length (*L*), the length of root that can ben freely in a pore. The free length promotes deflection, because the buckling force initiating the root bending inversely scales with the square of the pore size (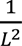 in equation 11, Supplementary Method, section 4).

The degree to which the base of the root is already embedded into the matrix also affects the ability of the root to penetrate through agarose gel cylinders. In our system, the principal effect was due to the support of the root base by the neighbouring cylinders reducing the pore size and therefore the free length *L*. It is possible that the compression of the root exerted by the lateral cylinders also limited the ability of the base of the root to rotate, but Euler’s buckling formula indicates that the prefactor *α* typically varies by a factor of 16 (from 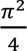 to 4*π*^2^) depending on how the root tip and base are fixed. With experiments showing that *α* was at most 4, potential variations in the buckling force due to the anchorage of the root base between agarose cylinders were less than a factor of 2. By comparison, penetrometer resistance of agricultural soils typically varies by a similar amount in response to changes in water content but can also vary by one order of magnitude in response to soil compaction (Gao *et al*., 2012). Root diameter can increase by more than 60% in response to soil compaction too (Dexter and Alston, 1991), which represents an increase of the moment of inertia of the root by a factor of 6.4. In our experimental system, we can conclude that anchorage at the base of the root was not the dominant factor. This may differ substantially under natural soil conditions, however, because our setup restricted observations to roots that were only weakly anchored between two cylinders.

The primary root trait used to overcome heterogeneities is the flexural rigidity of the root *E_r_I_r_*, with 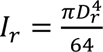, confirming the old studies of (Dexter and Hewitt, 1978; Whiteley *et al*., 1982; Whiteley and Dexter, 1984). The increase of the root diameters with soil strength (Iijima and Kato, 2007; Loades *et al*., 2013) has been interpreted as a mechanism to reduce axial stress and maintain the elongation rate in compacted soil, but experimental studies found weak correlations between root diameter and the ability to penetrate hard layers within genotypes from a diversity panel (Chimungu *et al*., 2015), thereby failing to support this hypothesis. Results presented here indicate a greater role of root diameter for preventing root deflection. Other traits that contribute to bending stiffness, such as root anatomy (Vanhees *et al*., 2020) and cell wall properties such as lignin or cellulose content (Kamchoom *et al*., 2023) may therefore represent valuable targets to adapt crops to soils’ mechanical limitations.

The coefficient of friction between root and soil (parameter *μ*), appeared to also have major effects on root deflections. The largest incidence angle above which penetration was no more possible (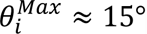) provided an indication of the magnitude of the root-agarose friction coefficient. Assuming Coulomb friction and the absence of bending moment or shear stress at the root tip, this incidence angle would mark the direction of growth at which slipping of the tip occurs, i.e. 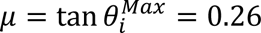. Because there are other forces resisting the deflection of the root due to the deformations at contact with the agarose cylinder, the friction angle is most likely lower than 15°. The value is similar to the characteristic angle of the distribution of incidence angle we measured (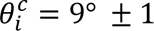, Figure 5B), which explains why deflection and penetration events were both frequently observed.

It is generally accepted that reduced root-soil friction is beneficial to growth (Mckenzie *et al*., 2013). Root developmental traits such as detachment of border cells (Bengough and Mckenzie, 1997) and the production of mucilage (Iijima *et al*., 2004; Vollsnes *et al*., 2010) are believed to help plant roots reduce friction. However, our results show that excessive friction reduction may increase the frequency of root deflections and result in more tortuous root systems. This may not be the case of all soils. Sand for example, which is more abrasive that agarose gel, is likely characterised by higher root-soil friction and is likely to prevent slippage (Bizet *et al*., 2016). Other soils however, such as soft clays (Miȩdlarz *et al*., 2019), are characterised by low friction angles and may be more prone to root deflections.

While root deflections events are often regarded as a negative consequence of growth in mechanically constraining soils (Tracy *et al*., 2012; Martins *et al*., 2020), their functional significance can only be fully understood when considered within a broader ecological context. Although root deflections increase horizontal dispersion of the root system (Dupuy *et al*., 2018), they also enable the root to minimise the mechanical energy required to deform the soil and thereby to sustain a larger elongation rate. The results presented here suggest that deflections are detrimental to the speed of the rooting front only when elongation rate within the obstacles is significantly larger than that in the pore space (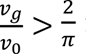 in our triangular system). Another important conclusion from this work is that the direction of deflection is not purely random, but instead shows a bias toward one randomly elective direction of growth. This observation, which mirrors patterns previously reported for hard obstacles (Yao *et al*., 2025), may also be indicative of a dispersion strategy, avoiding excessive accumulation of roots immediately below the stem.

**Table 1:**
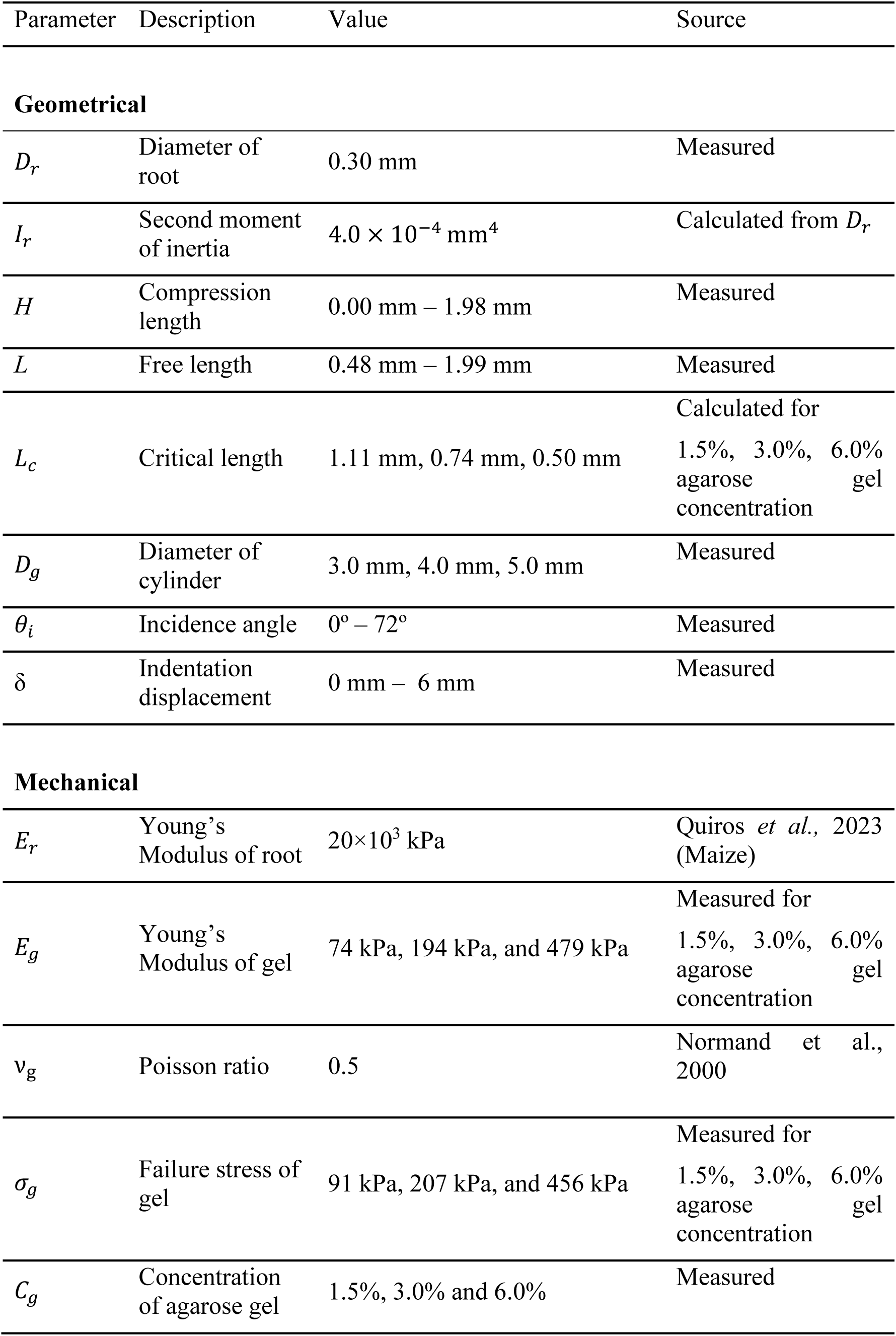
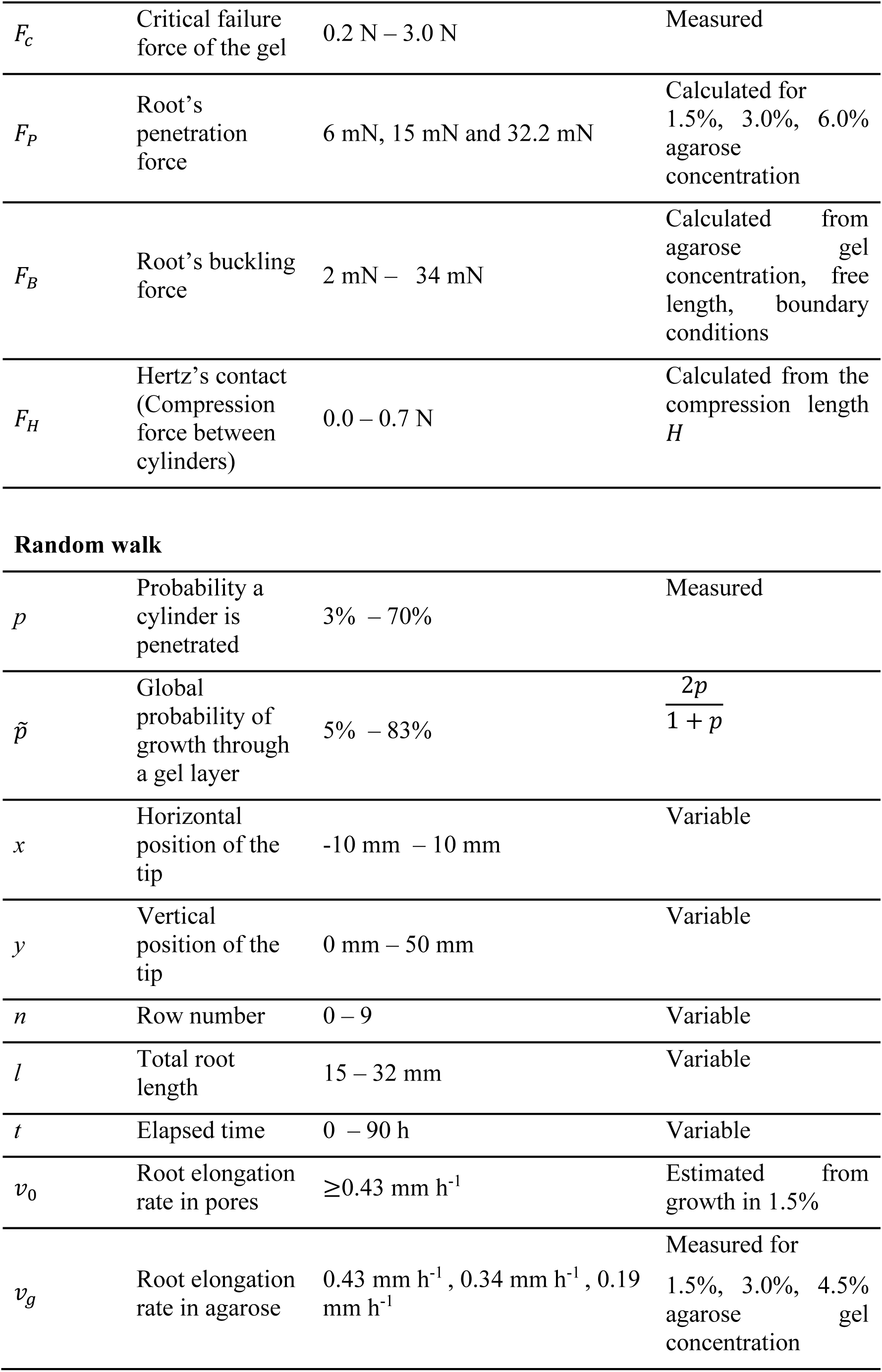
List of symbols and notations.

| Parameter | Description | Value | Source |
| --- | --- | --- | --- |
| **Geometrical** |  |  |  |
| $D_r$ | Diameter of root | 0.30 mm | Measured |
| $I_r$ | Second moment of inertia | $4.0 \times 10^{-4} \text{ mm}^4$ | Calculated from $D_r$ |
| $H$ | Compression length | 0.00 mm – 1.98 mm | Measured |
| $L$ | Free length | 0.48 mm – 1.99 mm | Measured |
| $L_c$ | Critical length | 1.11 mm, 0.74 mm, 0.50 mm | Calculated for 1.5%, 3.0%, 6.0% agarose gel concentration |
| $D_g$ | Diameter of cylinder | 3.0 mm, 4.0 mm, 5.0 mm | Measured |
| $\theta_i$ | Incidence angle | $0^\circ - 72^\circ$ | Measured |
| $\delta$ | Indentation displacement | 0 mm – 6 mm | Measured |
| **Mechanical** |  |  |  |
| $E_r$ | Young's Modulus of root | $20 \times 10^3 \text{ kPa}$ | Quiros *et al.*, 2023 (Maize) |
| $E_g$ | Young's Modulus of gel | 74 kPa, 194 kPa, and 479 kPa | Measured for 1.5%, 3.0%, 6.0% agarose gel concentration |
| $\nu_g$ | Poisson ratio | 0.5 | Normand *et al.*, 2000 |
| $\sigma_g$ | Failure stress of gel | 91 kPa, 207 kPa, and 456 kPa | Measured for 1.5%, 3.0%, 6.0% agarose gel concentration |
| $C_g$ | Concentration of agarose gel | 1.5%, 3.0% and 6.0% | Measured |
| $F_c$ | Critical failure force of the gel | 0.2 N – 3.0 N | Measured |
| $F_p$ | Root's penetration force | 6 mN, 15 mN and 32.2 mN | Calculated for 1.5%, 3.0%, 6.0% agarose concentration |
| $F_B$ | Root's buckling force | 2 mN – 34 mN | Calculated from agarose gel concentration, free length, boundary conditions |
| $F_H$ | Hertz's contact (Compression force between cylinders) | 0.0 – 0.7 N | Calculated from the compression length $H$ |
| Random walk |  |  |  |
| $p$ | Probability a cylinder is penetrated | 3% – 70% | Measured |
| $\tilde{p}$ | Global probability of growth through a gel layer | 5% – 83% | $\frac{2p}{1+p}$ |
| $x$ | Horizontal position of the tip | -10 mm – 10 mm | Variable |
| $y$ | Vertical position of the tip | 0 mm – 50 mm | Variable |
| $n$ | Row number | 0 – 9 | Variable |
| $l$ | Total root length | 15 – 32 mm | Variable |
| $t$ | Elapsed time | 0 – 90 h | Variable |
| $v_0$ | Root elongation rate in pores | $\geq 0.43 \text{ mm h}^{-1}$ | Estimated from growth in 1.5% |
| $v_g$ | Root elongation rate in agarose | $0.43 \text{ mm h}^{-1}$, $0.34 \text{ mm h}^{-1}$, $0.19 \text{ mm h}^{-1}$ | Measured for 1.5%, 3.0%, 4.5% agarose gel concentration |

## Acknowledgments

We thank Prof Jim Haseloff for advice on staining of agarose gel cylinders. The research was supported by the project BIOFLOW (PID2023-149435OR-I00) funded by MICIU/AEI /10.13039/501100011033 and by FEDER (EU). We thank the China Scholarship Council for funding the PhD of JY.

