## Supplementary Information for "Root responses to mechanical heterogeneity are shaped by pore geometry and root-soil biomechanics"

**SUPPLEMENTARY FIGURES**

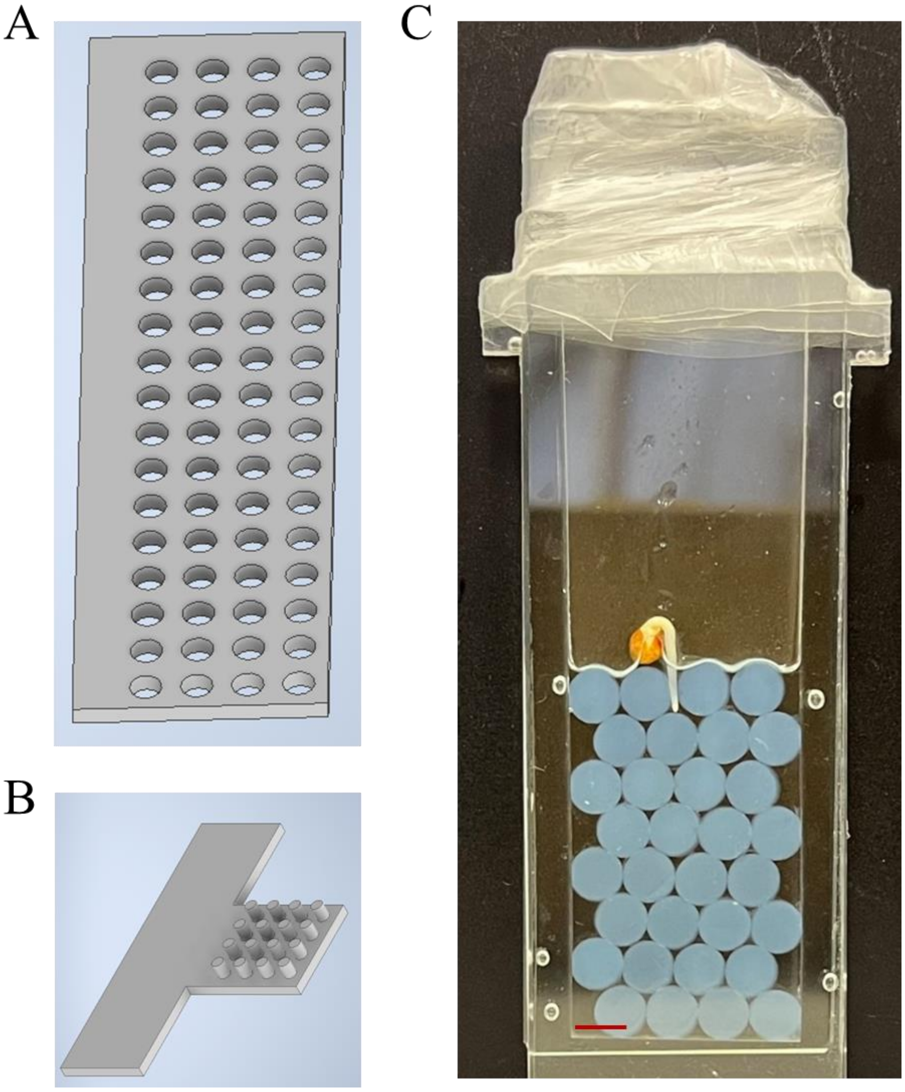


Figure S1. Preparation of agarose gel microcosm. (A) Moulds with holes of diameters of 3.0, 4.0 and 5.0 mm used for the fabrication of agarose gel cylinders. (B) Tool used for removing the agarose gel cylinders from the moulds contained pillars of diameters 2.5, 3.0 and 3.5 mm, slightly smaller than the corresponding hole diameters. (C) Example of a cabbage seedling planted in agarose gel cylinders of diameter 4.0 mm arranged in close packing inside a chamber made of a PDMS U-shape and two parallel glass slides. The red bar is 4 mm.

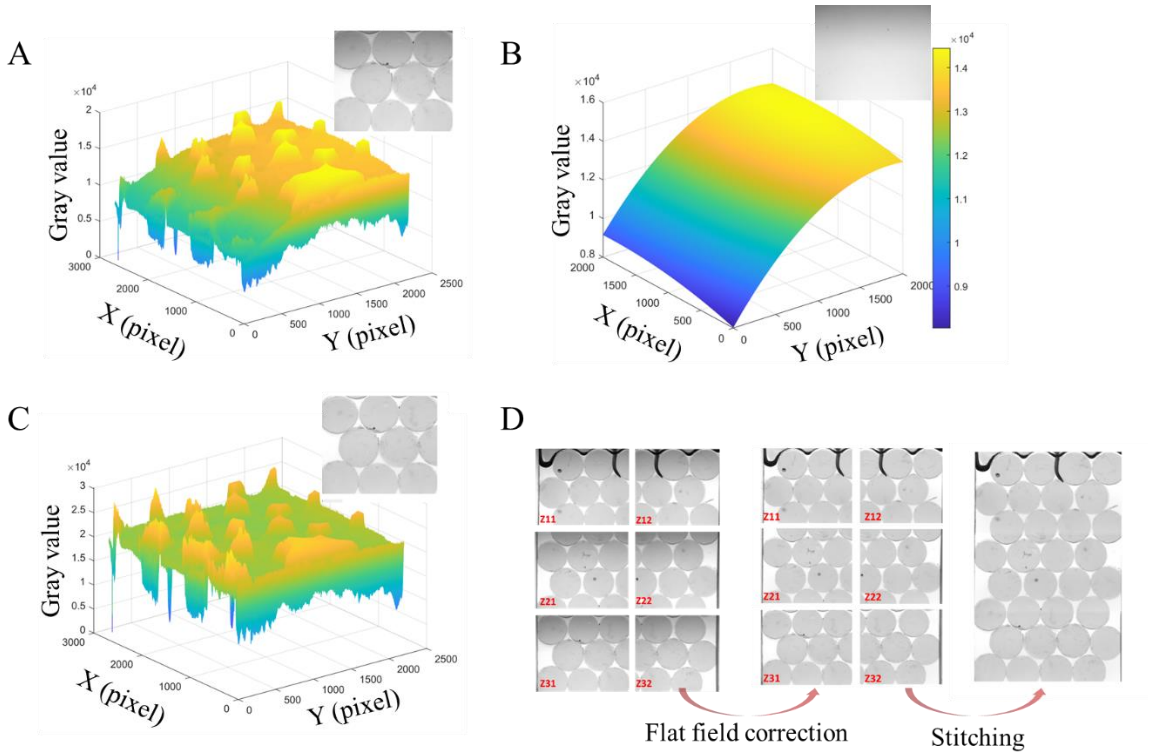


Figure S2. Flat field and stitching of image data. (A) Typical distribution of pixel intensity observed in the raw data when roots are present in the microcosm chamber containing granular substrate. The top right inset represents the raw image with agarose gel cylinders of 4.0 mm of diameter and at 3.0% agarose gel concentration. (B) Typical distribution of pixel intensity in the reference sample (here bulk agarose at a concentration of 3.0% which raw image is shown in the inset). (C) Distribution of pixel intensity after flat field correction and corresponding correction in the inset image. (D) Flat field correction and image reconstruction by stitching the 6 images.

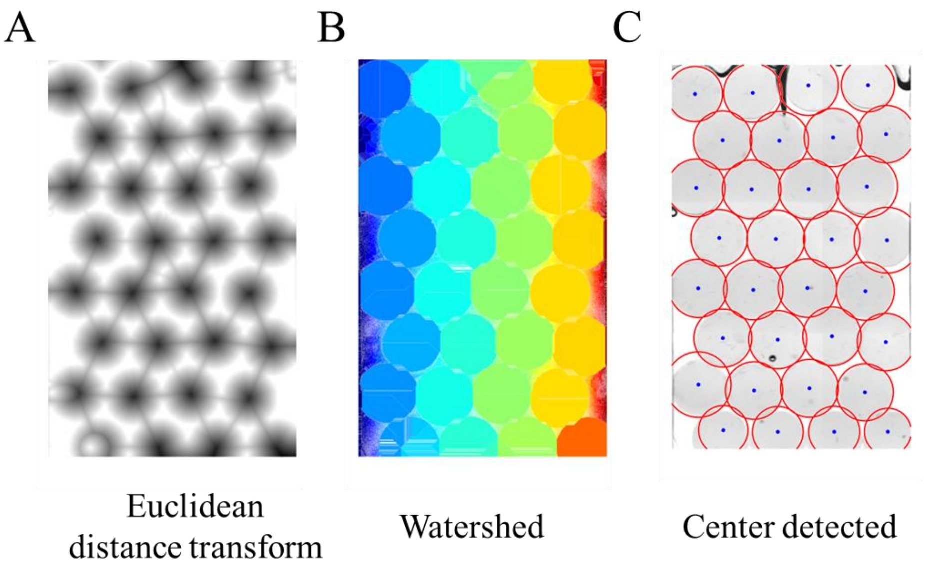


Figure S3. Detection of the centres of the agarose gel cylinders. (A) Euclidean distance transform was followed by (B) a watershed transform. (C) The centres of the cylinders are defined as the centres of mass of the segmented regions (blue points), with approximated boundaries (red circles) represented by circles with radius set as the major axis of the regionprops function from MATLAB.

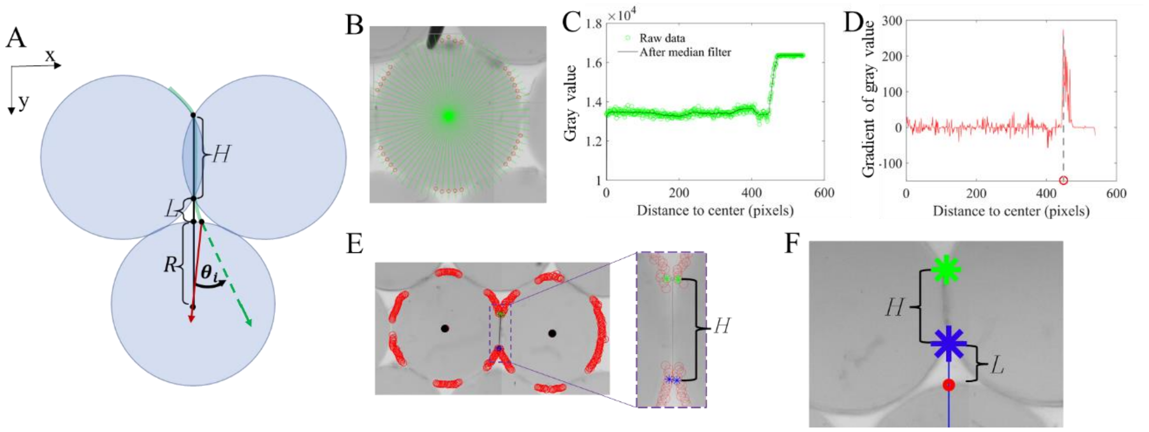


Figure S4. Extraction of parameters characterising the pore space when a root is interacting with a cylinder. (A) Sketch defining the compression length $H$, the free length $L$ and the incidence angle $\theta_i$ of the root tip on the cylinder. The continuous green curve symbolises the root skeleton. (B) Rays originating from the centre of the cylinder used to detect the boundary of the cylinder. (C) Pixel intensity profile along a ray and (D) the corresponding gradient along the profile. The position of the boundary point is indicated with the red circle. (E) Once boundary points have been detected along the circumference of the cylinder (left image of E), the compression length $H$ (right image of E) is defined from the pairs of nearest boundary points (green and blue asterisks) from two adjacent circles. (F) The free length is defined as the distance between the end point of the segment for compression length (blue asterisk) and the first point where the vertical ray intersects the next cylinder (red circle).

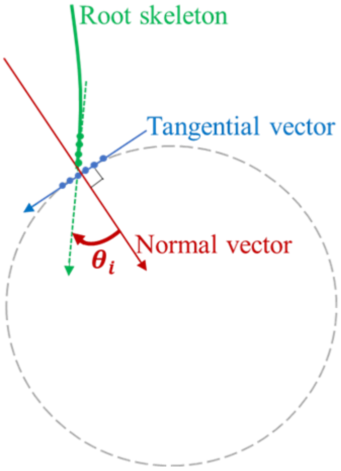


Figure S5. The incidence angle ($\theta_i$) is defined as the angle between the root tip vector (green dotted arrow) and the inward normal vector (red arrow) of the cylinder at the point of contact.

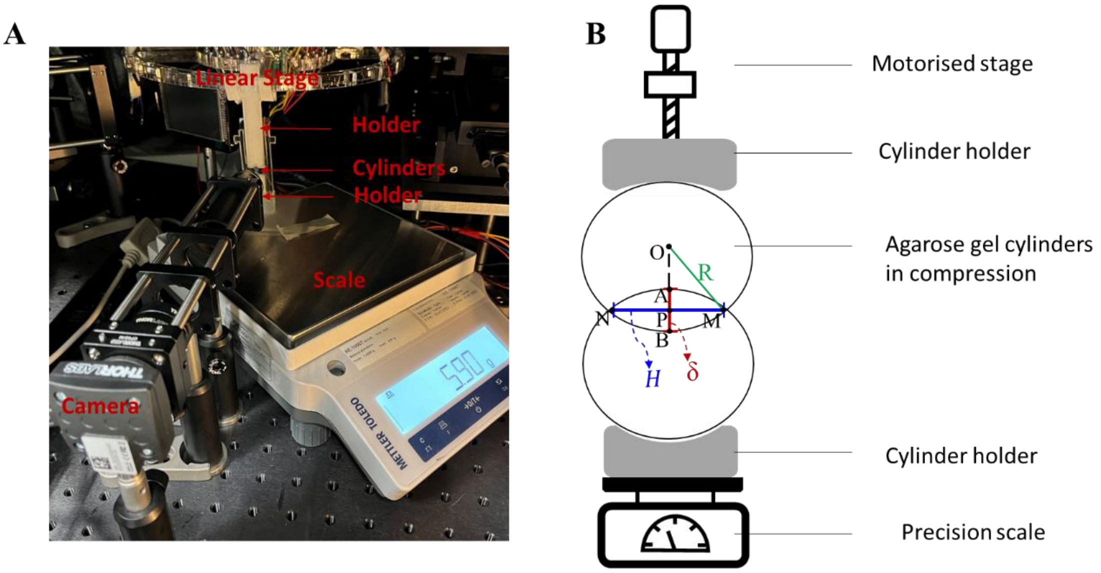


Figure S6. Compression test of agarose gel cylinders. (A) Experimental setup for the simultaneous acquisition of compression force $F_H$ and compression length $H$. (B) Schematics of the main parameters captured by a camera to test the validity of Hertz contact law between two cylinders.

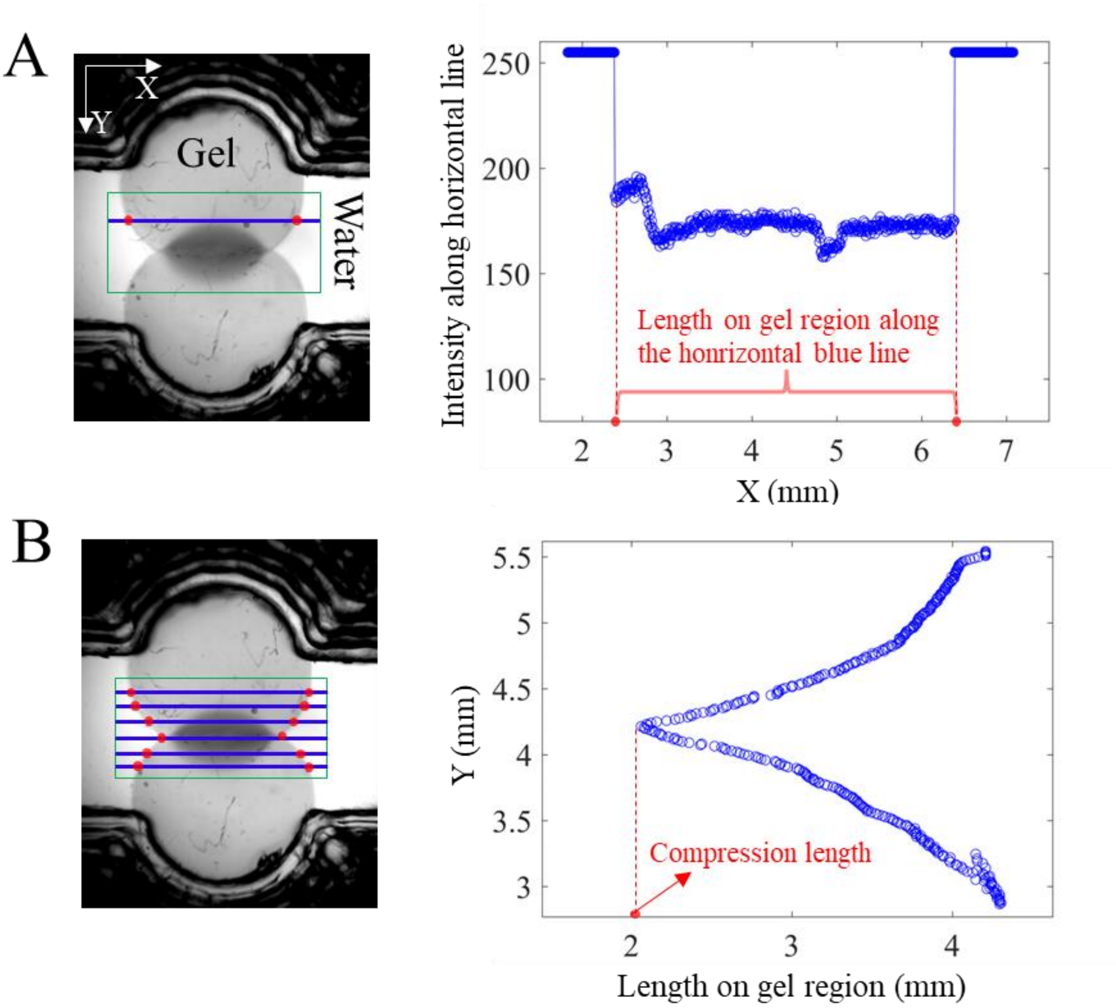


Figure S7. Determination of the compression length between two agarose gel cylinders during a compression test. (A) A green rectangular region is selected between the two cylinders, covering both the gel and the surrounding medium (water). This region defines the area where the algorithm is applied (left). The pixel intensity profile along a blue horizontal line (X axis) after contrast enhancement is shown on the right. Because the gel appears darker (lower intensity) than water, the length of the gel domain can be determined as the length of the unsaturated zone (red). (B) The length of the gel domain is determined for various horizontal lines identified by their Y-coordinate (left), and the compression length is determined as the smallest length of the gel domain (right).

### SUPPLEMENTARY METHOD

#### 1 – Staining the agarose gel

Reagents

- 1% periodic acid solution: 0.1 g periodic acid (P7875-25G, Merck, Spain) in 10 ml DI water
- Schiff reagent (10 ml): 0.19 g sodium metbisulfite
    - 0.3 ml 5N HCl
    - 9.7 ml DI water
    - It is better not to store longer than 2 weeks.
- Propidium iodide: 1 ml propidium iodide (P4864-10ML, Merck).

Staining the agarose gel cylinders

- Fabrication of the agarose gel cylinders using the protocol described in the main text (section *Preparation of the granular substrate* and Figure S1)
- Leave agarose cylinder in periodic acid solution for 30 minutes.
- Rinse thoroughly in DI water for 10 minutes
- Remove the water and add 1 ml Propidium iodide and 10 ml Schiff reagent solution, leave in for 2 hours, gel will be pink.
- Rinse thoroughly in DI water

Gel imaging

Light sheet fluorescence microscope (LSFM) was used for imaging the agarose gel cylinders (Liu *et al.*, 2021). Volume images were acquired with an excitation wavelength of 514 nm, and emission filters of 550 nm.

### 2 – Model of mechanical contact between two agarose gel cylinders

According to Hertz contact theory (P 63, Popov, 2010) with the geometrical parameters defined in Figure S6B, the force opposed to two parallel cylinders in contact with each other is related to the length of the cylinders $W$ and varies linearly with indentation $\delta$

$$F_H = \frac{\pi \tilde{E} W \delta}{4} \tag{1}$$

The indentation $\delta$ is related to $a$, the half of the contact width (here referred as compression length $H$),

$$\delta = \frac{a^2}{\tilde{R}} = \frac{H^2}{4\tilde{R}},$$

$\tilde{R}$ is the equivalent radius of the geometry of contact,

$$\frac{1}{\tilde{R}} = \frac{1}{R_1} + \frac{1}{R_2} = \frac{2}{R}. \tag{2}$$

so that the compression length $H$ is the related to the indentation

$$H = 2\sqrt{\tilde{R}\delta} = \sqrt{2R\delta} \tag{3}$$

and the force can be expressed as a function of the contact width $a$ (Johnson, 1985)

$$F_H = \frac{\pi (H/2)^2 \tilde{E} W}{4\tilde{R}}. \tag{4}$$

For two identical cylinders $\tilde{E}$ is defined by

$$\frac{1}{\tilde{E}} = \frac{1 - {\nu_1}^2}{E_1} + \frac{1 - {\nu_2}^2}{E_2} = 2\frac{1 - {\nu_g}^2}{E_g},$$

and $\nu_g$ is the Poisson ratio and $E_g$ is the Young's modulus of the gel. Replacing the value of $\tilde{E}$ and $\tilde{R}$ into equation (1) gives the relationships between the compression force $F_H$and the compression length $H$,

$$F_H = \frac{\pi E_g W H^2}{16(1 - {\nu_g}^2)R} \tag{5}$$

which is written non dimensionally as

$$\sqrt{\frac{8(1-\nu_g{}^2)F_H}{\pi W E_g D_g}} = \frac{2H}{D_g} \qquad (6)$$

with $D_g = 2R$ being the diameter of agarose cylinder.

Therefore, Hertz's theory for parallel cylinders predicts that the compression length $H$ is related to the square root of the compression force $F_H$.

#### 3 – Mechanical properties of bulk agarose and critical force required for penetration of agarose gel

The indentation tests of bulk agarose in Petri dishes showed a non-linear evolution of the force $F$ with the displacement of the translation stage, or equivalently with the indentation $\delta$ of the gel (Figure 1B). The force increase was steeper for higher concentrations of agarose. Tests carried out at different points on the gel surface followed the same curve of increasing force with indentation. However, the sudden collapse in force corresponding to the failure of the gel occurred at more variable indentation depths.

Hertz' framework was used to analyse the results of gel indentation tests. In the case of a rigid spherical indenter of radius $R_s$ in contact with an elastic infinite half space of Young's modulus $E_g$ and Poisson ratio $\nu_g$, the force resisting an indentation δ (Figure 1A) is

$$F = \frac{4}{3} \frac{E_g}{\left(1 - \nu_g{}^2\right)} R_s^{\frac{1}{2}} \, \delta^{\frac{3}{2}} \qquad (7)$$

This formula is well known in contact mechanics for elastic materials (Johnson, 1985), but not so well established for gels. Experimental results showed the relationship between the force $F$ and $R_s{}^{\frac{1}{2}}\delta^{\frac{3}{2}}$ (Figure 1C) was linear, which confirmed the validity of Hertz' framework (equation 7) for gels over the entire displacement range, even for large deformations just before failure.

During an indentation test, the force $F$, the radius of the spherical indenter $R_s$ and the magnitude of the indentation δ are known so that the Young's modulus of the gel $E_g$ can be determined if Poisson ratio is known. Agarose gel is a hydrogel, i.e. a polymer skeleton that traps water. We therefore hypothesized in a first approach that the agarose gel had the behaviour of an incompressible material, with a Poisson ratio of $\nu_g = 0.5$. We thus obtained:

$$F = \frac{16}{9} E_g R_s^{\frac{1}{2}} \delta^{\frac{3}{2}} \tag{8}$$

From the slope of $F$ versus $R_s^{\frac{1}{2}}\delta^{\frac{3}{2}}$, we obtained the Young's modulus $E_g$ for the different agarose concentrations (Figure 1D) with the smallest indenter radius (0.8 mm). As expected, the Young's modulus $E_g$ increases with the agarose concentration and the evolution is almost linear in the range of concentration. Note that the values we obtained are under the hypothesis of a Poisson ratio of 0.5 and might be overestimated in the very low-velocity regime where water has time to flow out the compressed zone (poroelastic effects).

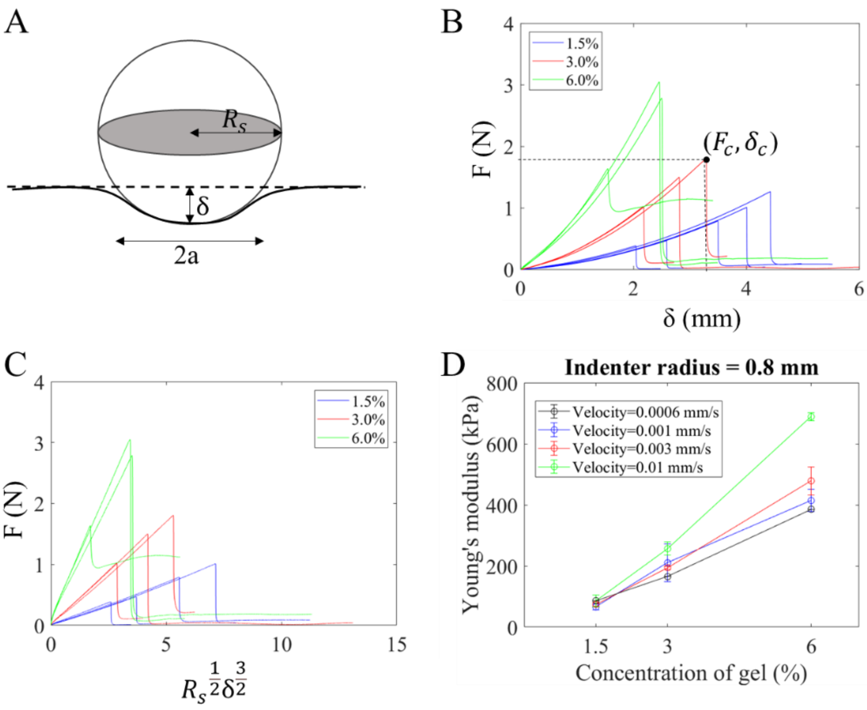


Figure 1. Mechanical properties of bulk agarose gel. (A) Definition of geometrical parameters used to apply Hertz' contact theory. (B) Force-displacement curves obtained at a displacement rate of 0.003 mm/s and with the spherical indenter

diameter ($2R_s$) of 1.6 mm (1/16 inch), with agarose gel concentration of 1.5 %, 3.0 % and 6.0 % (respectively blue, red and green curves). (C) Relationship between the indentation force and $R_s^{\frac{1}{2}}\delta^{\frac{3}{2}}$ confirms the validity of Hertz' contact theory. (D) Evolution of the Young's modulus with the concentration of agarose in the gel, for four different velocities of indentation and with $R_s$ =0.8 mm for the radius indenter.

When the stress reached a critical value, the gel failed and a catastrophic reduction of the force was observed. The peak force $F_C$ is the critical force required for the penetration of agarose gel. We recorded $F_C$ and the corresponding critical indentation depth $\delta_c$ (Figure 1B) during the indentation test. The failure stress $\sigma_\mathrm{g}$ is estimated from the peak force $F_C$ as:

$$\sigma_\mathrm{g} = \frac{F_C}{\pi a_c^2} \quad (9)$$

where $a_c$ is the radius of the contact area when the gel breaks. As we observed the Hertz's law to be valid up to failure, we deduced the value of $a_c$ from the indentation depth $\delta_c$ at failure, according to $a_c = \sqrt{R_s \delta_c}$. Therefore

$$\sigma_\mathrm{g} = \frac{F_C}{\pi R_s \delta_c} \quad (10)$$

### 4 – Mechanical model of root interaction with agarose gel cylinders

Under the effect of the root's axial growth against the agarose cylinder, the root is subjected to a compressive force. If the root behaves like a solid rod, when this force reaches the critical buckling force $F_B$, the root buckles and bends. The critical buckling force is characterised by the mechanical and geometric properties of the root. At first order, we assume the root apex part between cylinder rows $n-1$ and $n$ can be replaced by a cylindrical rod of constant diameter $D_r$ and uniform Young's modulus $E_r$ (Figure 2A). This root apex part experienced buckling for a critical force,

$$F_B = \alpha \frac{E_r\, I_r}{L^2}. \qquad (11)$$

Here, $E_r$ is the root's Young's modulus, $I_r$ is the second moment of inertia of the root calculated form the root's diameter $D_r$ with $I_r = \frac{\pi D_r^4}{64}$ for a plain cylinder, $L$ is the free length, i.e. the apex root length which is not supported laterally. The coefficient $\alpha$ is a dimensionless factor of order 1 which value depends on the boundary condition at the root tip and at its base and typically varies from $\pi^2/4$ to $4\pi^2$ .

The force $F_B$ must be compared to the penetration force $F_P$ needed for the root to penetrate the agarose gel. We assumed $F_P$ to be simply given by the failure stress of the gel, $\sigma_g$, depending on the agarose concentration, and by the cross-section of the root,

$$F_P = \frac{\sigma_g\, \pi D_r^2}{4}. \qquad (12)$$

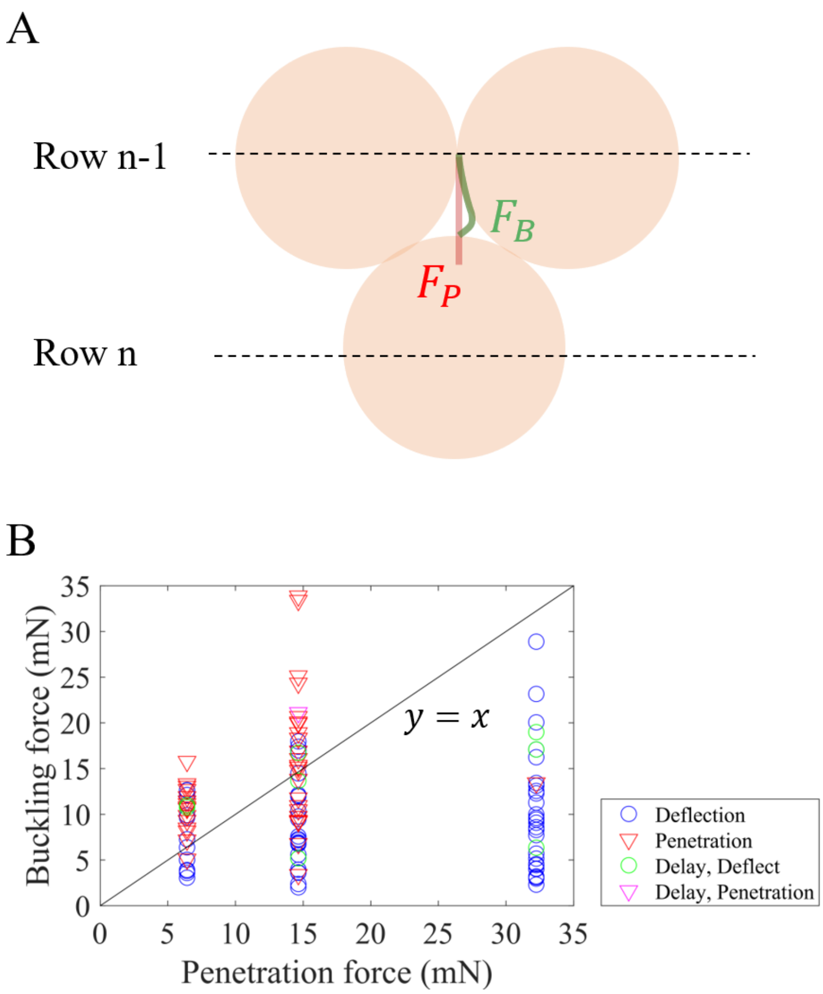


Figure 2. Penetration and buckling forces determine the root trajectory in the granular substrate. (A) Diagram showing the two ways a root can overcome a cylinder at row $n$. It can penetrate the gel if the penetration force $F_P$ is reached first, or deflect if the buckling force $F_B$ is reached first. (B) Buckling force $F_B$ calculated with $\alpha = 1$ vs Penetration force $F_P$. Colours indicate the root behaviours (red for penetration, blue for deflection, green for delay before deflection, magenta for delay before penetration). The black line corresponds to a slope of 1 with $F_B = F_P$.

The root is predicted to penetrate the gel when the penetration force is lower than the buckling force ($F_P < F_B$). If not ($F_B < F_P$), the root buckles before having reached the force necessary to break the gel. The criterium $F_B = F_P$ marks the transition between the two behaviors.

As a first approximation, we did not take into account the details of the boundary conditions and we set $\alpha = 1$. Potting $F_B(\alpha = 1) = \frac{E_r\, I_r}{L^2} = \frac{E_r \pi D_r^4}{64\, L^2}$ as a function of $F_P$ (Figure 2 B), gives quantitative indications on the validity of this approach. The plot shows that the values $F_B(\alpha = 1)$ and $F_P$ (5 to 35 mN) are of the same order of magnitude, which indicates mechanical buckling may be a good framework to analyse the experimental data. The criterium $F_B(\alpha = 1) = \frac{E_r\, I_r}{L^2} = F_P$ (line of slope 1) therefore provides an approximated rule to discriminate deflection cases from penetration cases.

Since $F_B$ is a function of the free length, the transition boundary $F_B = F_P$, can be expressed as a function of the critical length $L_c$ at which bending and buckling are equally likely to occur when $\alpha = 1$,

$$L_c = \sqrt{\frac{E_r D_r^2}{16\, \sigma_g}} = \frac{D_r}{4}\sqrt{\frac{E_r}{\sigma_g}}. \tag{13}$$

The buckling prefactor α characterising the environment of the growing root may be different from 1 depending on factors such as the anchorage at the base of the root at row $n - 1$, the pinning of the root at row $n$ or the incidence angle. At the transition boundary, the condition $F_B = F_P$ is equivalent to $\alpha \frac{{L_c}^2}{L^2} = 1$ or $\left(\frac{L}{L_c}\right)^2 = \alpha$. The dimensionless length ratio $(L/L_c)$ obtained at the transition boundary ($F_B = F_P$), provides a direct indication of the mechanical boundary conditions that the gel cylinders impose on the growing root.

### 5 - Random walk model of root growth in a heterogeneous substrate.

Here the objective is to characterise the effect of root deflections on root system properties that may affect the efficiency with which the plant can acquire soil resources (rooting depth, velocity of the root front, spread of the root distribution in soil). To achieve this objective, we propose models of root growth in the substrate that draws on concepts from random walk theory to describe the statistics of key root properties.

Experiments characterise the probability $p$ that a root interaction with an agarose gel cylinder results in a penetration event. However, a penetration event at row $n$ leads the root to grow straight up to row $n+2$ and skip the row $n+1$, whereas a deflection event causes the root to grow in the pore up to row $n+1$, where it interacts with a cylinder with probability of penetration $p$. To account for these different scenario, we chose to define the global probability $\tilde{p}$ that a root is growing straight from row $n$ to $n+1$. A root that grows through $m$ interactions with cylinders will grow through $n = 2pm + (1-p)m$ rows of cylinders. The global probability of growing through gel is $\tilde{p} = \frac{2pm}{2pm + (1-p)m} = \frac{2p}{1+p}$, and the probability of a root to grow in a pore is $1-\tilde{p}$. The values of $\tilde{p}$ computed from the experimental data of $p$ (presented in figures 4B and 4D of the main text) are given in table 1.

| Cylinders | $p$ | $\tilde{p}$ |
|---|---|---|
| **R=1.5 mm, C=3%** | 0.71 | 0.83 |
| **R=2.0 mm, C=3%** | 0.36 | 0.53 |
| **R=2.5 mm, C=3%** | 0.25 | 0.40 |
| **R=2.0 mm, C=1.5%** | 0.50 | 0.67 |

| | | |
|---|---|---|
| **R=2.0 mm, C=6%** | 0.03 | 0.06 |

Table 1. Values of global probability $\tilde{p}$ of growing through gel as a function of radius and gel concentration.

In the following steps, we consider a trinomial random walk with row increment $\Delta y = \frac{D_g\sqrt{3}}{2}$ due to the triangular packing of cylinders. Between 2 successive rows, we note $\Delta x$ the horizontal spread, $\Delta y$ the vertical increment and $\Delta l$ the growth increment, i.e. the root length increase. As the root encouters an agarose gel cylinder it can either penetrate the cylinder changing position and length by $\Delta x = 0, \Delta l = \Delta y = D_g\sqrt{3}/2$ at a growth velocity of $v_g$ with a probability of $\tilde{p}$. It can also deflect changing position and length by $\Delta x = \pm\frac{D_g}{2}, \Delta l = D_g\left(\sqrt{3} - 1 + \frac{\pi}{2}\right)/2$ at a growth velocity $v_0$ with a probability of $1 - \tilde{p}$ . In this case, we hypothesize the root tip experiences a vertical trajectory in the pore then advances by circling the cylinder in a quarter circle (Figure 3). There is equal probability $\frac{1-\tilde{p}}{2}$ of going left or right. The root may have different velocities when it grows impeded by the gel ($v_g$) or unimpeded when growing in the pore space ($v_0$). Therefore, the time needed to achieve a row increment is $\Delta t = \frac{D_g}{2}\left[\frac{(\sqrt{3}-1)}{v_0} + \frac{1}{v_g}\right]$ when the root is able to penetrate the agarose gel cylinder and $\Delta t = \frac{D_g}{2}\frac{\left(\sqrt{3}-1+\frac{\pi}{2}\right)}{v_0}$ when the root bypasses the agarose gel cylinder.

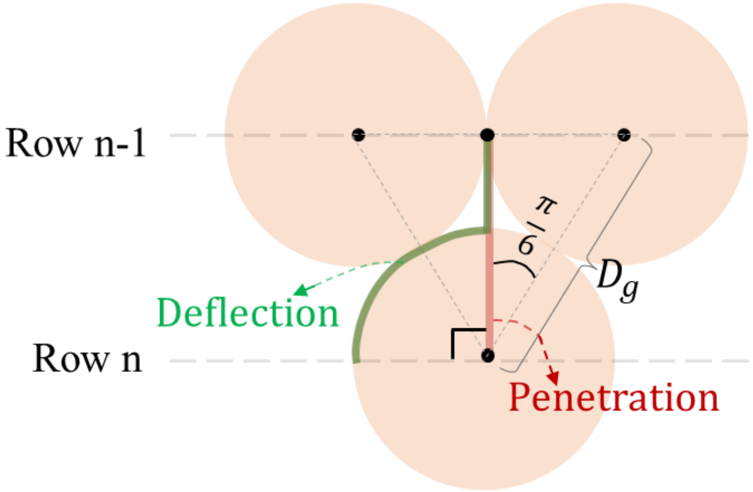


Figure 3. Schematic representation of the random walk model. The length needed to move from row $n-1$ to row $n$ depends on whether the root penetrates the gel (orange path) or whether it is deflected (green path). If the root penetrates the gel cylinder, the length of the root is the distance between the two rows, $l = D_g \frac{\sqrt{3}}{2}$. If the root is deflected, the length is calculated by considering the distance between row $n-1$ and the top of the cylinder at row $n$, that is, $D_g \left(\frac{\sqrt{3}-1}{2}\right)$, and then adding an additional one-quarter of the cylinder's circumference, that is, $\frac{D_g \pi}{4}$. Calculating the time to reach row $n$ is then obtained from the velocities of growth in the gel $v_g$ and the velocity of growth in the pore $v_0$.

After $n$ row increments, we consider the position of the root tip $(x, y)$ the length of the root $l$ and the time $t$,

$$x(n) = x(n-1) + \Delta x, \tag{14}$$

$$y(n) = y(n-1) + \frac{D_g\sqrt{3}}{2},$$

$$l(n) = l(n-1) + \Delta l,$$

$$t(n) = t(n-1) + \Delta t.$$

By taking into account the probabilities of each event, we get the expected values of increments:

$$\langle \Delta x \rangle = \tilde{p} \times 0 + \left(\frac{1-\tilde{p}}{2}\right)\frac{D_g}{2} - \left(\frac{1-\tilde{p}}{2}\right)\frac{D_g}{2} = 0,$$

$$\langle \Delta l \rangle = \tilde{p}\frac{D_g\sqrt{3}}{2} + (1-\tilde{p})\frac{D_g\left(\sqrt{3}-1+\frac{\pi}{2}\right)}{2}$$

$$\langle \Delta t \rangle = \tilde{p}\frac{D_g}{2}\left[\frac{\left(\sqrt{3}-1\right)}{v_0} + \frac{1}{v_g}\right] + (1-\tilde{p})\frac{D_g}{2}\frac{\left(\sqrt{3}-1+\frac{\pi}{2}\right)}{v_0}$$

The expected values of the root properties are therefore

$$\langle x(n) \rangle = \langle x(n-1) \rangle\,, \tag{15}$$

$$\langle y(n) \rangle = \langle y(n-1) \rangle + \frac{D_g\sqrt{3}}{2},$$

$$\langle l(n) \rangle = \langle l(n-1) \rangle + \frac{D_g}{2}\left[\left(\sqrt{3}-1+\frac{\pi}{2}\right) - \tilde{p}\left(\frac{\pi}{2}-1\right)\right],$$

$$\langle t(n) \rangle = \langle t(n-1) \rangle + \frac{D_g}{2}\left[\frac{\left(\sqrt{3}-1+\frac{\pi}{2}\right)}{v_0} - \tilde{p}\left(\frac{\pi}{2v_0} - \frac{1}{v_g}\right)\right].$$

And by recurrence

$$\langle x(n) \rangle = 0$$

$$\langle y(n) \rangle = \frac{nD_g\sqrt{3}}{2}$$

$$\langle l(n) \rangle = \frac{nD_g}{2}\left[\left(\sqrt{3}-1+\frac{\pi}{2}\right) - \tilde{p}\left(\frac{\pi}{2}-1\right)\right]$$

$$= \frac{\langle y(n) \rangle}{\sqrt{3}}\left[\left(\sqrt{3}-1+\frac{\pi}{2}\right) - \tilde{p}\left(\frac{\pi}{2}-1\right)\right]$$

$$\langle t(n) \rangle = \frac{nD_g}{2}\left[\frac{\left(\sqrt{3}-1+\frac{\pi}{2}\right)}{v_0} + \tilde{p}\left(\frac{1}{v_g} - \frac{\pi}{2v_0}\right)\right]$$

$$= \frac{\langle y(n) \rangle}{\sqrt{3}}\left[\frac{\left(\sqrt{3}-1+\frac{\pi}{2}\right)}{v_0} + \tilde{p}\left(\frac{1}{v_g} - \frac{\pi}{2v_0}\right)\right]$$

From these relationships, it can be concluded that the Length to Depth Ratio $LDR = \langle l(n)\rangle/\langle y(n)\rangle$, which inverse value indicates how efficiently plant roots progress vertically, decreases with the probability of penetration of the root. The LDR value is only dependent on the probability of penetration $\tilde{p}$, and it is only through that probability that the structure of the substrate affects the ability of the root to expand vertically (Figure 4D).

In terms of the time required for the root to reach a given depth, the model predicts that deflection is beneficial whenever the growth velocity inside the obstacle $v_g$ is too small and drops below a critical value,

$$v_g < \frac{2v_0}{\pi}. \tag{16}$$

The variance of the position of the root tip from the center of the row, $\langle x^2(n)\rangle$, is a good model indicator for a comparison with root growth patterns observed experimentally. The square of the distance $x$ at row $n$ is obtained as in equation 14,

$$x^2(n) = x^2(n-1) + 2\Delta x \times x(n-1) + \Delta x^2\ . \tag{17}$$

In the case where $\Delta x$ is independent of the previous steps, we have $\langle \Delta x \times x(n-1)\rangle = 0$ and the expected value is

$$\langle x^2(n)\rangle = \langle x^2(n-1)\rangle + \frac{(1-\tilde{p})D_g^2}{4} = \frac{n(1-\tilde{p}){D_g}^2}{4}. \tag{18}$$

To collapse the data obtained from the study and compare the data to model predictions, we use the following change of variable, $x = D_g x'$ and $n = 4n'/(1-\tilde{p})$. Using this change of variable, we transform equation 18 into a relationship independent of experimental parameters,

$$\langle x'^2(n')\rangle = n' \tag{19}$$

We compared model and observed values of $\langle x^2(n')\rangle$ using values of $\tilde{p}$ computed from measured $p$ (Table 1). Results showed the model systematically underestimates the variance of the distribution of the horizontal position of the root (Figure 4).

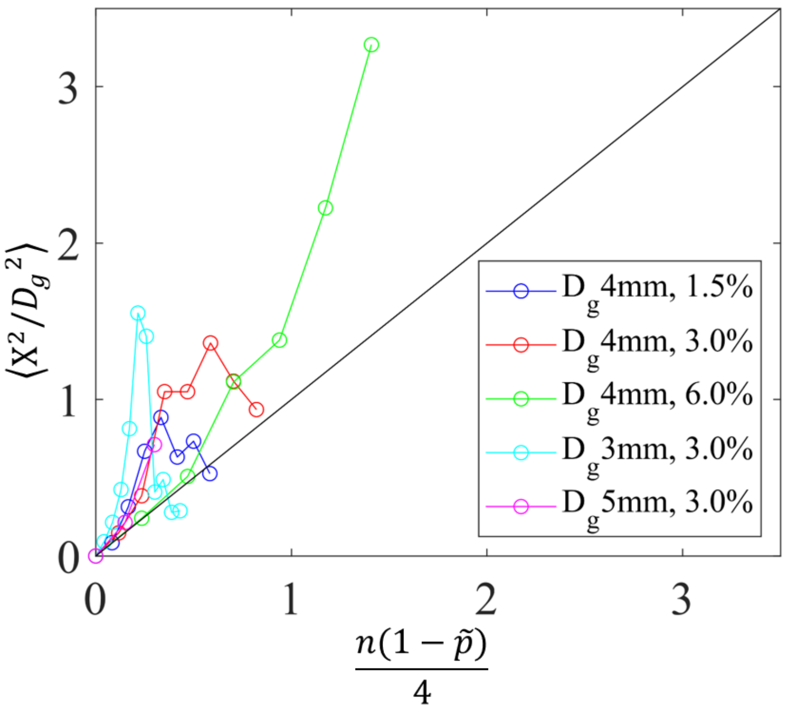


Figure 4: Collapse plot of the non dimensional variance of the horizontal position of the square of $X/D_g$ as a function of the normalised row number $n(1-\tilde{p})/4$. The model predicts equality of the two variables but experimental data were consistently larger than predicted.

Next, we considered whether the larger than expected values of $\langle x^2\rangle$ can be explained by a bias in the side of the deflection. If the root deflects always on the same side, following an oblique trajectory, the mean horizontal position is not null anymore,

$$\langle x(n)\rangle = \langle x(n-1)\rangle + \frac{(1-\tilde{p})D_g}{2} = n\frac{(1-\tilde{p})D_g}{2}$$

$x(n-1)$ and $\Delta x$ are still supposed to be independent variables but the expected value $\Delta x$ is strictly positive (if, for example, we systematically choose the rigth side for deflection) and equation 17 is transformed to

$$\langle x^2(n)\rangle = \langle x^2(n-1)\rangle + D_g\langle x(n-1)\rangle + \frac{(1-\tilde{p})D_g{}^2}{4}$$

and

$$\langle x^2(n)\rangle = D_g\sum_{i=1}^{i=n}\langle x(i-1)\rangle + n\frac{D_g{}^2(1-\tilde{p})}{4} = \frac{(1-\tilde{p})D_g^2}{2}\sum_{i=0}^{i=n-1} i + n\frac{(1-\tilde{p})D_g{}^2}{4}$$

$$\langle x^2(n)\rangle = \frac{(1-\tilde{p})D_g^2}{4}n(n-1) + n\frac{(1-\tilde{p})D_g{}^2}{4}$$

The expected value of $\langle x^2(n) \rangle$ is therefore

$$\langle x^2(n) \rangle = \frac{(1-\tilde{p})D_g^2}{4} n^2 \tag{20}$$
